# It runs in the family: Discovery of enzymes in the oleuropein pathway in Olive (*Olea europaea*) by comparative transcriptomics

**DOI:** 10.1101/2025.03.20.644408

**Authors:** Ornella Calderini, Mohamed O. Kamileen, Yoko Nakamura, Sarah Heinicke, Ryan M. Alam, Benke Hong, Yindi Jiang, Alma Gutiérrez, Fiammetta Alagna, Francesco Paolocci, Maria Cristina Valeri, Edoardo Franco, Soraya Mousavi, Roberto Mariotti, Lorenzo Caputi, Sarah E. O’Connor, Carlos E. Rodríguez-López

**Author notes:** to whom correspondence should be addressed: CERL,; SOC,; OC.

## Abstract

Olive (*Olea europaea* L.) is one of the most important crop trees, with olive oil being a key ingredient of the Mediterranean diet. Oleuropein, an oleoside-type secoiridoid, is the major determinant of flavor and quality of olive oil. Iridoid biosynthesis has been elucidated in *Catharanthus roseus*, which produces secologanin-type secoiridoids, but iridoid biosynthesis in other species remains unresolved. In this work, we sequenced RNA from olive fruit mesocarp of six commercial olive cultivars with varying oleuropein content, during maturation and ripening. Using this data we discovered three polyphenol oxidases with oleuropein synthase (OS) activity, a novel oleoside-11-methyl ester glucosyl transferase (OMEGT) synthesizing a potential intermediate in the route, and a 7-*epi*-loganic acid O-methyltransferase (7eLAMT). Interestingly, integrating transcriptomics data from 15 plant species from three iridoid-producing plant orders (Lamiales, Gentianales, and Cornales), and tissue expression panels from *Jasminum sambac* and *Fraxinus excelsior*, we discovered two 2-oxoglutarate dependent dioxygenases (named 7eLAS) that synthesize 7-*epi*-loganic acid; in contrast *C. roseus* 7-deoxy-loganic acid hydroxylase (7DLH), a known bottleneck in MIA production, is a cytochrome p450. This comparative co-expression method, which combines guilt by association and comparative transcriptomics approaches, can successfully leverage big datasets for untargeted discovery of enzymes.

**Key Findings:**

- Expression of genes involved in iridoid biosynthesis, from the early MEP pathway to the last step of oleuropein biosynthesis, decreases during olive fruit maturation.
- We discovered an oxoglutarate dependent dioxygenase, 7-*epi*-loganic acid synthase (7eLAS), catalyzing the stereoselective oxidation of 7-deoxy-loganic acid to 7-*epi*-loganic acid, in a reaction analogous to *C. roseus* 7-deoxy-loganic acid hydroxylase (7DLH), a cytochrome p450.
- We report a 7-*epi*-loganic acid O-methyltransferase (7eLAMT) orthologous to *Catharanthus roseus* loganic acid O-methyltransferase and found a novel oleoside-11-methyl ester glucosyl transferase (OMEGT) synthesizing 7-β-1-D-glucopyranosyl-oleoside-11-methyl ester, a potential intermediate in the oleuropein biosynthesis route.
- We discovered three olive polyphenol oxidases that have oleuropein synthase (OS) activity, catalyzing the conversion of ligstroside to oleuropein.

## INTRODUCTION

Olive (*Olea europaea* L.) is one of the most culturally important crops of the Middle Eastern and Mediterranean cultures. Olive oil has been an important part of the Mediterranean diet for millennia, to such an extent that the word for oil in most European languages is derived from the word for olive (Hoad 2003; “Aceite.,” Real Academia Española). One of the major components of olive oil, and a determinant of flavor and quality, is oleuropein: a phenolic secoiridoid ester that composes 6-14% of the dry weight of the fruit (Amiot, Fleuriet, and Macheix 1986; Ryan, Robards, and Lavee 1999). Previous research has found that genes involved in oleuropein biosynthesis, namely iridoid synthase (ISY) and oleoside-11-methyl ester (OME) synthase, are more highly expressed in domesticated than in wild olives (Rodríguez-López et al. 2021) highlighting the importance of the secoiridoid biosynthetic pathway.

Iridoid glycosides, non-canonical monoterpenes characterized by a cyclopentanopyran fused ring, are one of the most widespread specialized metabolites, reported to be present across the largest group of flowering plants, the Asterids (Stull et al. 2018). Despite their abundance, iridoids remain critically understudied, and most of what we know of their biosynthesis has been elucidated in the “non-model model” *Catharanthus roseus* (Apocynaceae.) Oleuropein is derived from the oleoside-type secoiridoid OME, which differs from the secologanin-type iridoids (e.g., iridoids present in *C. roseus*) by a characteristic exocyclic olefin (**Figure 1**) and has been reported to be present across the Oleaceae with varying concentrations and derivatives (Jensen, Franzyk, and Wallander 2002). Despite their importance, their biosynthetic pathway has not been fully elucidated, though the early steps are assumed to be the same as in the well-studied *C. roseus*.

**Figure 1.**
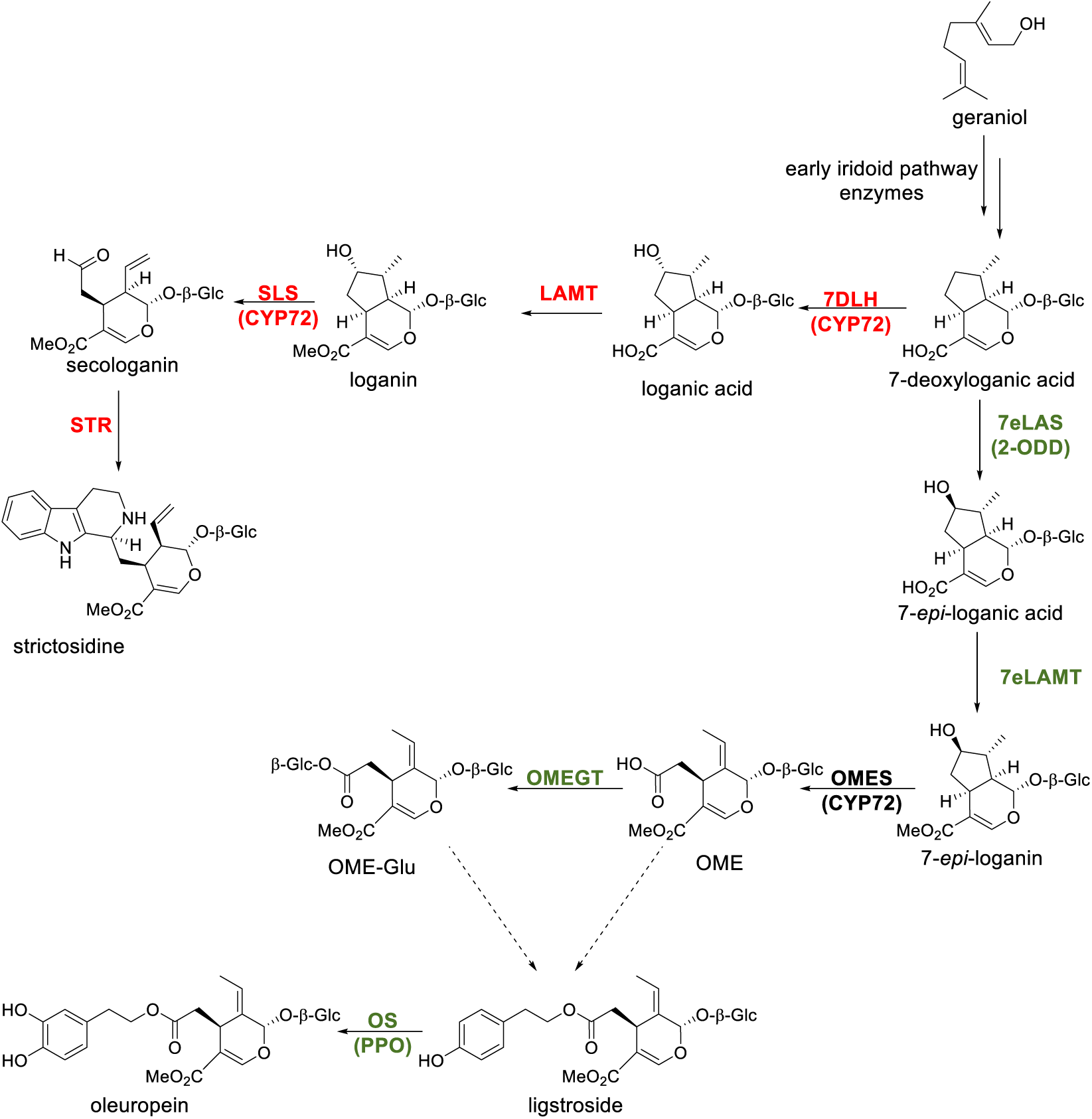
Secoiridoid biosynthetic pathways in three commonly studied species. Enzymes marked in red have been characterized in *Catharanthus roseus* and in black we show previously characterized enzymes in *Olea europaea*; in olive green we show enzymes characterized in this work. Enzyme abbreviations: 7DLH, 7-deoxy-loganic acid hydroxylase; 7eLAS, 7-*epi*-loganic acid synthase; SLS, secologanin synthase; STR, strictosidine synthase; LAMT, loganic acid O-methyltransferase; 7eLAMT, 7-*epi*-loganic acid O-methyltransferase; OMES, oleoside methyl ester synthase; OMEGT, oleoside-11-methyl ester glucosyl transferase; OS, oleuropein synthase. Protein families: CYP72, cytochrome p450 CYP72 family; 2-ODD, 2-oxoglutarate-dependent dioxygenase; PPO, polyphenol oxidase. Compounds: OME, oleoside-11-methyl ester; OME-Glc, 7-β-1-D-glucopyranosyl oleoside-11-methyl ester.

In this work, we sequenced RNA from olive fruit mesocarp of six commercial olive cultivars with varying oleuropein accumulation, during maturation and ripening. Using this data, we performed a comparative co-expression approach, integrating transcriptomics data from 15 plant species from three iridoid-producing plant families (Lamiales, Gentianales, and Cornales), including tissue expression panels from two oleoside-type iridoid producing species (*Jasminum sambac* and *Fraxinus excelsior*). With this approach we discovered two oxoglutarate dependent dioxygenases that produce 7-*epi*-loganic acid in a stereoselective manner, that we named 7-*epi*-loganic acid synthase (7eLAS) to differentiate from *C. roseus* 7-deoxy-loganic acid hydroxylase (7DLH), a cytochrome p450. Using homology-based approaches we also report the discovery of 7-*epi*-loganic acid O-methyltransferase (7eLAMT) and the novel oleoside-11-methyl ester glucosyl transferase (OMEGT), completing the pathway to 7-β-1-D-glucopyranosyl-oleoside-11-methyl ester, reconstructed in *Nicotiana benthamiana*. Finally, we discovered two enzymes, belonging to the polyphenol oxidase family, with Oleuropein Synthase (OS) activity, producing oleuropein when incubated with ligstroside. This work sets a precedent for leveraging publicly available datasets through a comparative approach, robustly narrowing down gene candidates from tens of thousands to a few hundred, allowing broader hypotheses on the nature of the enzyme candidates.

## METHODS

### Iridoid standards

Loganin, secologanin, ligstroside, oleuropein and salidroside standards were purchased from Sigma-Aldrich; 8-*epi*-loganin from AnalytiCon Discovery GmbH (Potsdam, Germany) and OME was purchased from PhytoLab GmbH (Vestenbergsgreuth, Germany). The rest of the iridoid standards (7-deoxy-loganic acid, 7-deoxy-8-*epi*-loganic acid, 7-deoxy-loganin, 7-deoxy-8-*epi*-loganic acid, 7-*epi*-loganic acid, 7-*epi*-loganin, and 7-ketologanin) were synthesized from commercially available geniposide (*Biosynth Carbosynth*, Staad, Switzerland) as previously reported (Rodríguez-López et al. 2021).

### Plant material

Olives from Moraiolo, Coratina, Leccino, Arbequina, Tendellone, Dolce d’Andria cultivars were harvested from collection fields of CNR IBBR in Perugia (Italy) every 20 days from 45 (stage 1) until 125 (stage 5) days after flowering (DAF). Olive fruit mesocarp was frozen in liquid nitrogen, pulverized using a mortar and pestle, and stored at – 80 °C until needed.

### RNA-Seq analysis

RNA was extracted from frozen and milled olive fruit mesocarp using Plant RNeasy® Kits (QIAGEN; Hilden, Germany) and sent for RNA-Sequencing to BGI Genomics (Shenzhen, China) following the company’s protocols, which included mRNA enrichment, library preparation and paired-end sequencing (2x150.) Namely: raw reads quality was assessed using fastQC (v0.11.5; (“FastQC” 2016)), results aggregated using multiQC (v1.17;(Ewels et al. 2016)), and processed using trimmomatic (Bolger, Lohse, and Usadel 2014). Trimmed files were mapped against the published genomes Farga (Cruz et al. 2016) and Arbequina (Rao et al. 2021) olive cultivars using HISAT2 (v2.2.1; (Kim et al. 2019)). Genome guided transcriptome assembly was performed using Trinity (v2.8.5; (B. J. Haas et al. 2013; Grabherr et al. 2011)) with a max intron length of 2500, resulting contigs were cleaned from duplicates using CDHIT (v4.7; (Fu et al. 2012)) with a 90% identity threshold, keeping the longest contig, and coding sequences were predicted using TransDecoder (B. Haas, n.d.). Genes were annotated by taking the predicted protein sequences and running them through eggNOG mapper (v2.1.12-1; (Cantalapiedra et al. 2021; Huerta-Cepas et al. 2019)) using an E-value cutoff of 1x10^-3^, and threshold score of 60, 40% identity and 20% coverage, and using DIAMOND (Buchfink, Reuter, and Drost 2021). Expression was estimated by quasi-mapping the runs using salmon (v0.14.1;(Patro et al. 2017)).

The genome, gene models and predicted peptide sequences were obtained from published work for *Antirrhinum majus* (M. Li et al. 2019; Tavares et al. 2018), *Camptotheca acuminata* (Zhao et al. 2017), *Callicarpa americana* (Hamilton et al. 2020), *Catharanthus roseus* (C. Li et al. 2023), *Cinchona pubescens* (Nataly Allasi et al. 2022), *Gelsemium sempervirens* (Franke et al. 2019), *Mitragyna speciosa* (Brose et al. 2021), *Rauvolfia tetraphylla* (Stander et al. 2023) and *Sesamum indicum* (Wang et al. 2022). For the rest of the species, only the genome was used, with no annotation, and the same pipeline as stated above was followed to generate a genome-guided transcriptome, the difference being that raw sequencing data was obtained from the Sequence Read Archive, with project numbers corresponding to each species as shown **Supplementary Table 1**. The genomes used for these species were obtained from GenBank Assemblies (GA) and annotated using the bioproject (BP) as follows: *Forsythia suspensa* (GA: GCA_023638005.1; BP: PRJNA793127), *Fraxinus excelsior* (GCA_019097785.1; BP: PRJEB4958), *Jasminum sambac* (GA: GCA_018223645.1; BP: PRJNA723725), *Osmanthus fragrans* (GA: GCA_019395295.1; PRJNA529305) and *Penstemon barbatus* (GA: GCA_003313485.2; BP: PRJNA479669). Only for *J. sambac* and *F. excelsior*, expression was estimated by quasi-mapping, using salmon (v0.14.1; (Patro et al. 2017)). Orthogroup inference was performed using OrthoFinder (v2.5.5;(Emms and Kelly 2019)) integrating the predicted peptides of all of the above mentioned species.

### Cloning methods

RNA was extracted from Olive fruits using Plant RNeasy® Kits (QIAGEN; Hilden, Germany), and cDNA libraries prepared using the SuperScript IV VILO MM (ThermoFisher; Waltham, United States). Candidate genes were amplified using Platinum SuperFi PCR MM (ThermoFisher; Waltham, United States) and cloned using ClonExpress II (Vazyme; Nanjing, China) into either pOPINF (OMEGT) or pOPINM (7eLAMT) for heterologous expression in *E. coli*, or directly into 3Ω1 destination vector for heterologous expression in *Nicotiana benthamiana*. Plasmids were propagated in *E. coli* Top10, and sequences confirmed by Sanger sequencing (GENEWIZ Germany GmbH, Leipzig, Germany).

### Reconstitution in Nicotiana benthamiana

Sequence-verified plasmids were transformed into *Agrobacterium tumefaciens* GV3101 via electroporation, plated on LB agar with antibiotics (50 μg·ml^-1^ rifampicin, 50 μg·ml^-1^ gentamicin and 200 μg·ml^-1^ spectinomycin) and single colonies picked and confirmed by colony PCR using Phire HotStart II Master Mix (ThermoFisher; Waltham, United States). Positive colonies were inoculated in liquid LB with the above-mentioned antibiotics, grown overnight at 28 °C and 220 rpm agitation in the dark, and pelleted by centrifugation at 5000 RCF for 5 minutes. The pellet was resuspended in as much infiltration buffer (50 mM MES pH 5.5, 10 mM MgCl_2_, and 200 μM acetosyringone) as needed to reach an optical density (OD_600_) of 0.6 and incubated in darkness at 28 °C and 220 rpm for 2 hours. When more than one gene was to be infiltrated, an equivolumetric mixture was done previous to incubation. Three weeks old *Nicotiana benthamiana* plants were selected, and the abaxial side of selected leaves was infiltrated with the Agrobacterium solution, using a needleless syringe. After 72 hours, substrate was infiltrated in infiltration buffer without acetosyringone, and 96 hours later, infiltrated tissue was isolated, frozen in liquid Nitrogen, and extracted with 10 volumes of methanol. The extract was sonicated, centrifuged and filtered through a 0.45 μm PTFE filter, and injected to the HPLC for analysis.

### Heterologous expression in *Escherichia coli*

For protein production in *E. coli*, sequence-verified plasmids were transformed into *E. coli* BL21 via heat shock transformation, plated on LB with carbenicillin (100 μg·ml^-1^) and single colonies confirmed by PCR, as stated above. Positive individual colonies were dropped in 5ml liquid LB broth with carbenicillin and incubated overnight at 37 °C and 220 rpm. This pre-inoculum was then added to 100 ml fresh YT media with carbenicillin and incubated at 37 °C and 220 rpm and transferred to 18 °C and 220 rpm when an OD_600_ of 0.6 was reached. IPTG was added to a final concentration of 0.5 mM to induce protein expression and incubated overnight. Cells were harvested by centrifugation (5000 RCF for 10 minutes), the pellet was weighted, and protein was extracted by using the B-PER cell lysis kit (ThermoFisher; Waltham, United States) following manufacturer instructions. His-tagged proteins were purified from the clarified solution by incubating with 100 μl of NiNTA-agarose beads (Qiagen) for 1 hour at 4 °C, and eluted using B1 buffer (50 mM Tris-HCl, 50 mM glycine, 500 mM NaCl, and 250 mM imidazole, pH 8.0). Buffer was exchanged to A4 buffer (20 mM HEPES, 150 mM NaCl and 10% glycerol, pH 7.5) by serial dilutions, using Amicon 30-kDa size-exclusion concentrators (Millipore.) Protein concentration was estimated by measuring absorbance at 280 nm in a nanodrop (ThermoFisher) and estimating the molar extinction coefficient based on the protein sequence, using Expasy ProtParam (Gasteiger et al. 2005).

### Enzymatic assays *in vitro*

To assay for methyltransferase activity, purified enzyme was diluted to a final concentration 0.1 g·l^-1^ and reactions were performed in 50 μl of 50 mM Tris buffer (pH 8) with freshly added 0.1% (v/v) β-mercaptoethanol, 50 μM ascorbic acid, and 100 μM *S*-adenosyl methionine. The reaction was started by adding 50 μM substrate, and stopped after 2 hours by adding 100 μl ice-cold methanol.

To assay for methyltransferase activity, purified enzyme was diluted to a final concentration 0.1 g·l^-1^ and reactions were performed in 50 μl of 50mM Tris buffer (pH 8) with freshly added 0.1% (v/v) β-mercaptoethanol, 50 μM ascorbic acid, and 100 μM S-adenosyl methionine (SAM.) For assaying UDP-glucosyl transferase activity, 0.1 g·l^-1^ enzyme was incubated with 250 μM UDP-glucose in a 50 mM Tris buffer (pH 7.5). For the assays to obtain the pH optima, the same conditions were kept but used a 50 mM MES buffer for lower pH. Reactions were started by adding 50 μM substrate, incubated at 30 °C, and stopped after 2 hours by adding 100 μl ice-cold methanol. Assays were centrifuged and filtered through a 0.45 μm PTFE filter previous to HPLC injection.

### Identification of 7-β-1-D-glucopyranosyl oleoside-11-methyl ester (OME-Glc)

His-tagged enzyme was produced in *E. coli* in a large, 1 l batch, and purified using Ni-agarose columns in an ÄKTA^TM^ FPLC system (Cytiva). A large-volume reaction was performed by setting 100 parallel 200 μl reactions, 50 mM MES at pH 5. and was subjected to semi-preparative high-performance liquid chromatography (HPLC) for product isolation. An Agilent 1260 Infinity II HPLC instrument connected to an autosampler, diode array detector (DAD), and fraction collector for compound detection and isolation. Chromatographic separation was performed using a Phenomenex Kinetex XB-C18 (5.0 μm, 100 Å, 100 × 2.1 mm) column maintained at 40 °C under gradient elution using reversed phase conditions. The mobile phases used for separation were water with 0.1% formic acid (A) and acetonitrile (B). The flow rate was set at 1.5 ml·min^-1^, and chromatographic separation was performed at 5% B for 2 min, followed by a linear gradient from 5% to 10% B in 12 min, 90% B for 3 min, 10% B for 3 min (t_total_ 20 min). Prior to injection, the samples were diluted to 1 mg·ml^-1^ with methanol and filtered using a 0.22 µm PTFE syringe filter. The diluted samples were placed in the autosampler, 20 µl injections were performed, and fractions were collected by monitoring the UV 254 nm and 238 nm. Fractions were pooled and evaporated to dryness. The isolated compound was then submitted for NMR analysis.

### NMR Characterization

NMR spectra of enzymatically generated 7-glucopyranosyl oleoside-11-methyl ester were measured by a 700 MHz Bruker Avance III HD spectrometer (Bruker Biospin GmbH, Rheinstetten, Germany), equipped with a TCI cryoprobe using standard pulse sequences as implemented in Bruker Topspin ver. 3.6.1. (Bruker Biospin GmbH, Rheinstetten, Germany) at 298 K. Chemical shifts were referenced to the residual solvent signals of MeOH-*d_3_* (*δ*_H_ 3.31/*δ*_C_ 49.0). The assignment and spectra are shown in **Supplementary Figures 1** through **7**. The chemical shifts agreed with the published data (Kuwajima et al. 1989).

### Metabolite profiling using HPLC-MS

Metabolite profiling was performed as previously stated (Rodríguez-López et al. 2021) with minor modifications. Namely, samples were chromatographically separated using a Thermo UltiMate 3000 UHPLC system (Thermo Fisher Scientific) equipped with an Acquity UPLC BEH C18 column (2.1 × 50 mm, 1.7 µm, 100 Å; Waters), coupled via pneumatic-assisted ESI to an Impact II q-TOF mass spectrometer (Bruker Daltonik). Iridoids were separated at 40 °C using a gradient from 0.1% formic acid in water to acetonitrile, following the previously reported gradients (Rodríguez-López et al. 2021; 2022). The output was ionized in negative mode, with a capillary voltage of 3.5 kV and a nebulizer pressure of 2.5 bar, with nitrogen as drying gas (flow of 11 l·min^-1^, 350 °C.) Data dependent fragmentation was triggered on an absolute threshold of 400 and acquired on the most intense peaks, excluded after 3 events, with dynamic collision energy from 20 to 50 eV. Raw MS files were converted to mzXML using Bruker Data Analysis software (Bruker Daltonik, Bremen, Germany), and, when needed, extracted ion chromatograms (XICs) were exported to csv using MZmine2 v2.40.1 (Pluskal et al. 2010).

### Statistics

Unless otherwise specified, data analysis was performed using the *base* library of the R programming language (v 4.4.1, R Core Team). Figures were generated with the aid of the *ggplot2* (Wickham 2016), *gplots* (Warnes et al. 2009) and *pheatmap* (Kolde 2019) libraries, and Venn diagrams were produced using the *VennDiagram* (Chen and Boutros 2011) package. Differential expression analysis, including count normalization, was performed using the Likelihood Ratio Test option from *DESeq2* (Love, Huber, and Anders 2014) and self-organizing maps were done via the *kohonen* library (Wehrens and Kruisselbrink 2018). The package *seqinr* (Charif and Lobry 2007) was used to handle nucleotide and peptide sequences and *ape* (Paradis, Claude, and Strimmer 2004) was used for tree handling and phylogenetic analyses. The tree of UDP-glucose transferases (UGT) was generated using a multiple sequence alignment using MUSCLE (Edgar 2004; Madeira et al. 2024) and the tree were was generated using ModelFinder (Kalyaanamoorthy et al. 2017) via IQ-TREE (Nguyen et al. 2015), and the LAMT orthogroup tree was selected from Orthofinder standard output results; both trees were plotted using iTOL v6 (Letunic and Bork 2024). Extracted ion chromatograms were extracted from raw files using MZmine2 (Pluskal et al. 2010), were plotted using Excel® and molecules drawn using ChemDraw®.

## RESULTS

### Transcriptomic profiling of olive fruit mesocarp through maturation and ripening

We collected fruits of six commercial olive cultivars for sequencing. The sweet varieties *Dolce d’Andria* and *Tendellone,* varieties used to produce table olives because of low oleuropein levels, as well as four varieties containing medium to high levels of phenolic secoiridoids: *Arbequina, Leccino*, *Coratina* and *Moraiolo* (Mousavi et al. 2022; Alagna et al. 2012). We analyzed the content of oleuropein and oleuropein precursors through maturation and ripening at five different time points. With the exception of oleoside-11-methyl ester (OME), all measured metabolites decreased as the fruit matured in the tree, in accordance with the available literature (**Figure 2**) (reviewed by Skodra et al. 2021.) Interestingly, varieties with low to moderate iridoid levels have an initial oleuropein content similar to high oleuropein cultivars, suggesting that oleuropein degradation plays a role in oleuropein levels of maturing fruit (**Supplementary Figure 8**.)

**Figure 2.**
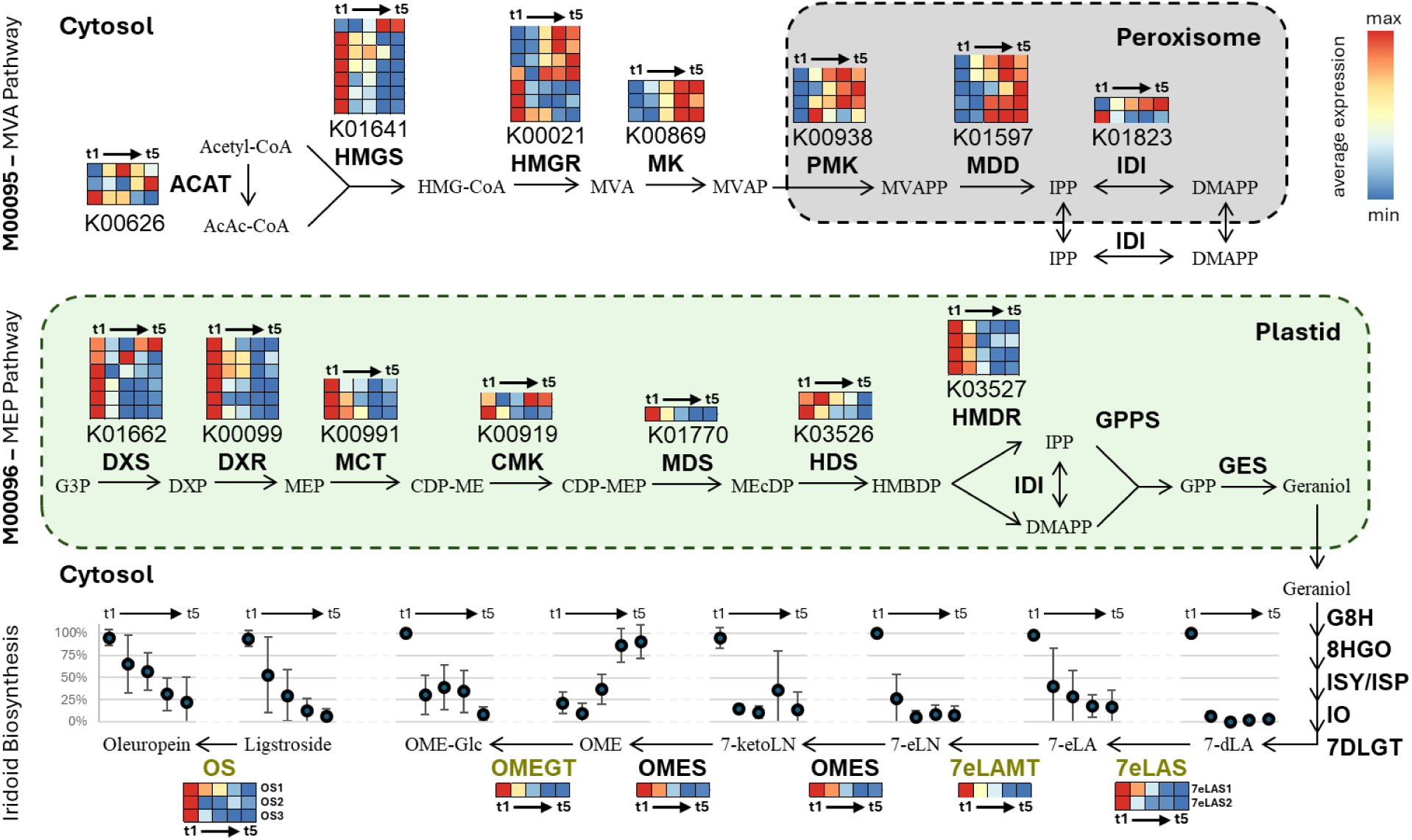
Enriched KEGG modules and iridoid biosynthetic genes. Schematic of differentially expressed genes (DEG) belonging to enriched KEGG modules of terpenoid biosynthesis, increasing (M00095 – MVA pathway; top) or decreasing (M00096 – MEP pathway; center) during olive maturation and ripening. The secoiridoid biosynthetic pathway is shown at the bottom, with enzymes characterized in this work shown in olive green; ligstroside biosynthesis remains unsolved and thus it is represented as disconnected. Heatmaps show expression patterns of all DEGs (one per row) annotated as the KEGG orthology number shown below each plot, with colors showing the average of the log-transformed transcripts per million (log_2_(TPM+1)) of all species, with color-scale ranging from the minimum (blue) to the maximum (red) mean expression value per gene (row.) Scatter plots show the mean (dots) and standard deviation (bars) of the intensity, scaled by species, of standard-confirmed iridoids through the five measured stages of maturation. MVA pathway: ACAT, acetyl coenzyme A (CoA) acetyltransferase; HMGS, hydroxymethylglutaryl-CoA synthase; HMGR, hydroxymethylglutaryl-CoA reductase; MK, mevalonate kinase; PMK, phosphomevalonate kinase; MDD, mevalonate diphosphate decarboxylase; isopentenyl diphosphate delta-isomerase (IDI). MEP pathway: DXS, 1-deoxy-D-xylulose-5-phosphate synthase; DXR, 1-deoxy-D-xylulose-5-phosphate reductoisomerase;MCT, 2-C-methyl-D-erythritol 4-phosphate cytidylyltransferase; CMK, 4-(cytidine 5′-diphospho)-2-C-methyl-D-erythritol kinase; MDS, 2-C-methyl-D-erythritol 2,4-cyclodiphosphate synthase; HDS, 4-hydroxy-3-methylbut-2-en-1-yl diphosphate synthase; HMDR, 1-hydroxy-2-methyl-2-(E)-butenyl 4-diphosphate reductase; GPPS, geranyl pyrophosphate synthase. Iridoid biosynthesis: GES, geraniol synthase; G8H, geraniol 8-hydroxylase; 8HGO, 8-hydroxygeraniol oxidoreductase; ISY, iridoid synthase; ISP, iridoid synthase paralogue; IO, iridoid oxidase; 7DLGT, 7-deoxyloganetic acid glucosyltransferase; 7eLAS, 7-*epi*-loganic acid synthase; 7eLAMT, 7-*epi*-loganic acid O-methyltransferase; OMES, oleoside methyl ester synthase; OMEGT, oleoside-11-methyl ester glucosyl transferase; OS, oleuropein synthase.

Using the same tissue from which metabolites were extracted, we performed an RNA-Seq experiment and assembled a genome-guided transcriptome using the published Farga genome (Cruz et al. 2016) (**Supplementary Figure 9A**.) The resulting transcriptome assembly had an ExN50 of > 2000 kb at 90% of expression and a mapping average of 90% (**Supplementary Figure 9B**.) PCA analysis showed that most changes in expression were due to maturation, with the main principal component (PC1), explaining 24% of the variance, clearly separating the samples by maturation stage (**Supplementary Figure 10**.) Interestingly, although PC2 (8% of the variance) separates samples by cultivar within each ripening state (**Supplementary Figure 10**), no separation was consistent with ligstroside or oleuropein content in the first 11 components, with a cumulative 75% of explained variance.

A likelihood ratio test, blocked by cultivar, revealed a total of 41,182 differentially expressed genes (DEGs, FDR < 0.01) changing during ripening. Using a self-organizing map (SOM) for dimensionality reduction and applying hierarchical clustering analysis (HCA) on the resulting codebook vectors, DEGs were clustered into eight distinct type expression patterns (**Supplementary Figure 11**), falling into two basic categories: increasing (**Supplementary Figure 12A-D**) or decreasing (**Supplementary Figure 12E-H**) with maturation, at different rates. An enrichment analysis shows that genes upregulated as olive fruit matures are enriched in KEGG pathway annotations consistent with sugar catabolism and respiration as well as fatty acid biosynthesis (**Supplementary Table 2**.) On the other hand, downregulated genes are overrepresented in processes related to photosynthesis, cell wall, and biosynthesis of secondary metabolites, particularly terpenoids (**Supplementary Table 2**.) Interestingly, enrichment analysis of KEGG Module annotations suggest that the mevalonate pathway is upregulated, while the non-mevalonate pathway is downregulated as maturation progresses (**Figure 2; Supplementary Table 3**). Iridoids are derived from geraniol produced by the MEP pathway (Contin et al. 1998). Consistently, the accumulation of measured metabolites (with the exception of oleoside methyl ester) and the expression pattern of secoiridoid biosynthetic genes decreases through ripening (**Figure 2**.) Thus, for discovering the missing enzymes in the oleuropein biosynthetic pathway, we will focus on the 24,857 genes differentially downregulated through ripening.

### Comparative co-expression analysis reveals that an oxoglutarate dependent dioxygenase hydroxylates 7-deoxy-loganic acid to produce 7-*epi*-loganic acid

The early steps of iridoid biosynthesis are the same in both olive and *C. roseus*. The first step in which the chemistry diverges is after the formation of 7-deoxy-loganic acid. In *C. roseus*, 7-deoxy-loganic acid is hydroxylated by 7DLH (Cr7DLH), a cytochrome p450 from the CYP72 family, to form loganic acid (Salim et al. 2013), while in olive, 7-deoxy-loganic acid is hydroxylated to form 7-*epi*-loganic acid. We initially assumed that the 7-*epi*-loganic acid synthase (7eLAS) from olive would be a homologue of Cr7DLH; nonetheless, no olive protein with sequence similarity to Cr7DLH showed any hydroxylation activity on 7-deoxy-loganic acid. We thus widened our search; however, a guilt by association approach yielded too many gene candidates to test, due to the confounding factor of fruit ripening.

Since the early iridoid pathway is shared across the Asterids, we performed a comparative co-expression analysis to reduce the number of candidate genes. Genome guided transcriptome assemblies were generated for five Lamiales species and integrated with published data from other iridoid producers within the Asterids, for a total of 15 plant species: nine Lamiales, five Gentianales, and one Cornales (**Supplementary Table 1**). We focused on members of the Oleaceae within our selection that have reliable reports of secoiridoid accumulation, and available RNA-Seq data of aerial, underground and reproductive tissues. Thus, we selected project PRJNA723725, having RNASeq data on leaf, flower, stem and root tissues of *Jasminum sambac*; and project PRJEB4958, which contains RNASeq data on leaf, flower, cambium and root tissues of *Fraxinus excelsior.* It has been reported that *J. sambac* and *F. excelsior* accumulate secoiridoids in leaf tissue (Damtoft, Franzyk, and Jensen 1992; Jensen, Franzyk, and Wallander 2002; Ross et al. 1982) and although no information is available for other tissues, we can reasonably assume that at least one of the selected tissues will have a little to no secoiridoid biosynthesis. RNA-Seq experiments were mapped against their respective genome guided assemblies, and expression patterns estimated using a *z*-score calculated on the log-transformed transcripts per million (TPM). The codebook vectors from SOMs were used to perform an HCA, that allowed us to visually distinguish eight type expression patterns (**Figure 3**.) For each species, we selected the best BLAST results of biosynthetic genes in the early iridoid pathway against each genome guided assembly, and we selected the cluster that contained the highest count of early biosynthesis gene candidates as the cluster likely to contain the missing step (**Supplementary Figures 13** and 14).

**Figure 3.**
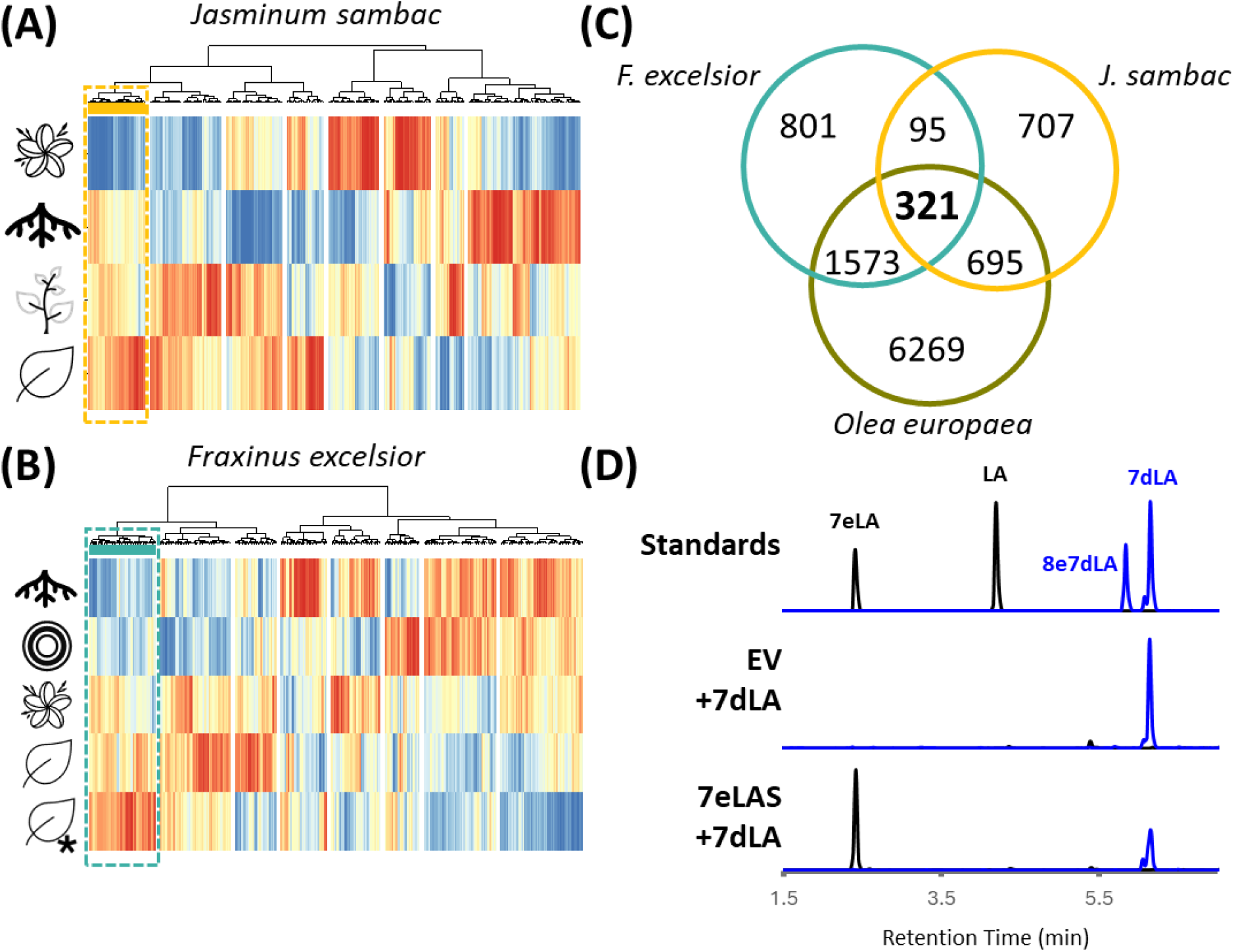
Comparative transcriptomics of Oleaceae species. Heatmap of the 400 self-organizing map nodes showing the type expression patterns of *Jasminum sambac* (A) and *Fraxinus excelsior* (B) in different tissues, denoted by icons; the selected clusters, in yellow and turquoise, contain candidate biosynthetic genes in the early iridoids pathway. From top to bottom: flower, root, stem, and leaves of *J. sambac*, and root, cambium (concentric circles), flower, leaf and leaf of a selfed tree (leaf + asterisk) of *F. excelsior*. (C) Venn diagram of orthogroup membership of genes belonging to the clustersof *J. sambac*, and *F. excelsior*, and the differentially expressed genes from *Olea europaea* that decrease during ripening; a total of 321 orthogroups are shared among the three gene selections. (D) Extracted **i**on **c**hromatogram (XIC) of the most abundant adducts of loganic acid (LA, [M-H]^-^) and 7-*epi*-loganic acid (7eLA, [M-H]^-^) in black (375.1297 ± 0.05); and 7-deoxy-loganic acid (7dLA, [M-H]^-^) and 8-*epi*-7-deoxy-loganic acid (7dLA, [M-H]^-^) in blue (359.1348 ± 0.05). From top to bottom: mix of standards (Standards), and extracts of *N. benthamiana* leaves co-infiltrated with 7-deoxy-loganic acid and Agrobacterium carrying an empty vector (EV+7dLA) or 7-*epi*-loganic acid synthase (7eLAS+7dLA.)

As seen in **Supplementary Figure 13**, in *J. sambac* candidates of Iridoid Synthase Paralogue (ISP), Iridoid Oxidase (IO), 7-deoxyloganetic acid glucosyl transferase (DLGT) and OMES group together, in a cluster that is expressed in every tissue except flower (**Figure 3A**); we thus selected 2,561 transcripts (belonging to 1,818 orthogroups) with that expression pattern for further analysis. Interestingly, in *F. excelsior*, candidate biosynthetic genes (ISY, ISP, IO, 7eLAMT, and OMES) also cluster together (**Supplementary Figure 14**), but their expression patterns show expression in leaf and flower, and little to no expression in root and cambium (**Figure 3B**), yielding almost twice as many candidates (4,238 genes; 2,790 orthogroups). These candidate lists were integrated with the DEG from olive ripening, only keeping the genes that were differentially expressed in olive during ripening, that had an orthogroup member in the candidate list for *J. sambac* and *F. excelsior*. There were 321 orthogroups that met these criteria (**Figure 3C**), which led to a reduction of olive gene candidates from the initial 24,857 DEG to 789, from which only 332 had a Pfam annotation.

From these candidates, annotations related to oxidases include eight multicopper oxidases, unlikely to catalyze the expected reaction and not explored further, six cytochrome p450 enzymes, five oxoglutarate dependent dioxygenases (ODDs), one Rieske oxygenase and one peroxidase. Testing via transient expression in *N. benthamiana* leaves, infiltrating 7-deoxy-loganic acid as a substrate, contigs TRINITY_GG_13709_c0_g1_i1 and TRINITY_GG_36519_c0_g1_i2, annotated as ODDs, were shown to consume 7-deoxy-loganic acid and produce 7-*epi*-loganic acid in a stereoselective manner, with no other epimer being detected (**Figure 3D**). To avoid confusion with the 7DLH reported in *C. roseus*, which is a cytochrome p450, these ODDs were named *O. europaea* 7-*epi*-loganic acid synthase one (Oe7eLAS 1) and two (Oe7eLAS 2), respectively. Interestingly, despite having low co-expression with each other (r = 0.46), coding sequences have 91% identity and peptide sequences share a 95% similarity. Notably, *C. roseus* has eight members in this orthogroup, one being deacetoxyvindoline 4-hydroxylase (D4H), an enzyme from the late MIA pathway expressed in leaf idioblasts (C. Li et al. 2023).

### 7-epi-loganic acid O-methylransferase (7eLAMT)

After formation of 7-*epi*-loganic acid, a methyltransferase is predicted to convert this intermediate to 7-*epi*-loganin. Analyzing the resulting orthogroups, we extracted the sequences belonging to *C. roseus* LAMT (CrLAMT) orthogroup (OG0000240), which catalyzes the formation of loganin from loganic acid. It can be observed from a phylogenetic analysis (**Figure 4**) that olive orthologues of interest cluster within the Lamiales species, in a clade adjacent to the Gentianales, where CrLAMT is located. The contig TRINITY_GG_16319_c0_g1_i4 from the *O. europaea* assembly clustered with the Lamiales LAMTs, appeared to be a full-length protein, and was differentially expressed during maturation, so it was selected as a likely pathway candidate. When tested *in vitro*, the heterologously expressed, purified protein consumed 7-*epi*-loganic acid to produce 7-*epi*-loganin (**Figure 4**) and was thus named *O. europaea* 7-*epi*-loganic acid O-methyltransferase (Oe7eLAMT.) Interestingly, the enzyme shows no measurable activity when fed 7-deoxy-loganic acid (**Figure 4**), which points at a route analogous to *C. roseus* where 7-deoxy-loganic acid is first oxidized and then methoxylated (**Figure 1**.)

**Figure 4.**
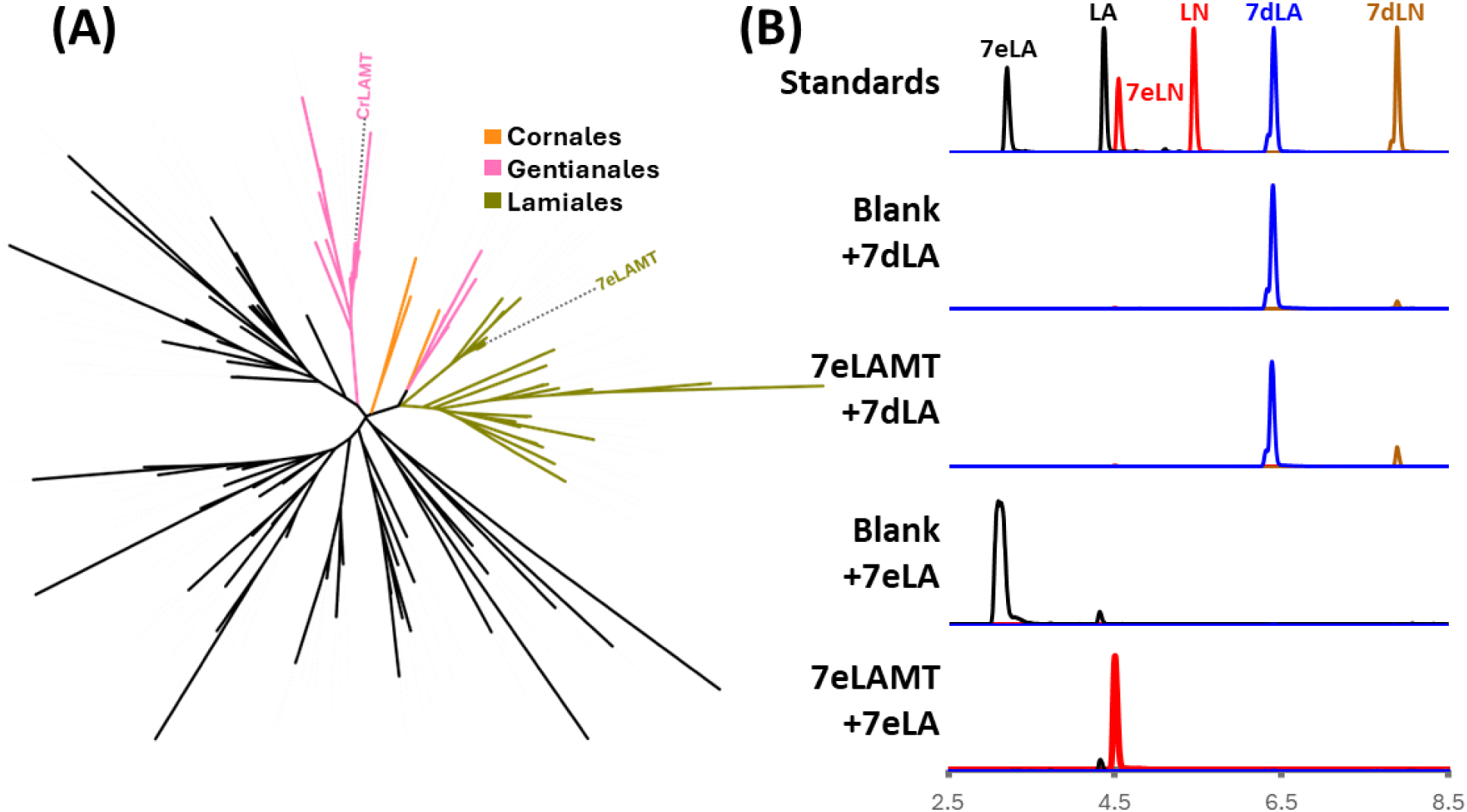
Characterization of 7-*epi*-loganic acid O-methyltransferase (Oe7eLAMT.) (A) Unrooted Orthofinder tree of group OG0000240, including orthologues of *Catharanthus roseus* loganic acid O-methyltransferase (CrLAMT) from 15 plant species. The lades of the closest orthologues are highlighted according to their corresponding order: Lamiales (olive green), Gentianales (pink) and Cornales (orange.) (B) Extracted ion chromatogram (XIC) of the most abundant adducts of loganic acid (LA, [M-H]^-^) and 7-*epi*-loganic acid (7eLA, [M-H]^-^) in black (375.1297 ± 0.05); loganin (LN, [M+formate]^-^) and 7-*epi*-loganin (7eLN, [M+formate]^-^) in red (435.1508 ± 0.05); 7-deoxy-loganic acid (7dLA, [M-H]^-^) in blue (359.1348 ± 0.05); and 7-deoxy-loganin (7dLN, [M+formate]^-^) in brown (419.1559 ± 0.05). From top to bottom: mix of standards (Standards), negative control and purified protein reactions incubated with 7-deoxy-loganic acid (Blank+7dLA and 7eLAMT+7dLA, respectively) and with 7-*epi*-loganic acid (Blank+7eLA and 7eLAMT+7eLA.)

### OME glucosyl transferase (OMEGT)

After formation of 7-*epi*-loganin, the previously reported CYP72 OMES catalyzes the formation of OME, which is then converted through an unknown mechanism to ligstroside (**Figure 1**). Analysis of the LC-MS profiling of olive fruits revealed a chromatographic peak which had a high correlation with 7-*epi*-loganin and secoxyloganin, and a fragmentation pattern matching an iridoid having two hexoses, matching literature reports of a glycoside moiety of oleoside-11-methyl ester (Kuwajima et al. 1989). We speculated that a glycosylated OME product may be an on-pathway intermediate to oleuropein. To identify a biosynthetic gene that is responsible for this glucosylation, we gathered the genes having a Pfam annotation of UDP glucosyl transferases (UDPGT) among differentially expressed genes in fruit. After phylogenetic analysis of these sequences (**Figure 5**), we searched for UGTs that are part of phylogenetic group L (AtUGT75B1/B2), enzymes that are known to recognize carboxylic groups and catalyze glucose ester bond formation (Caputi et al. 2012). Candidate TRINITY_GG_29808_c0_g1_i1 was selected for heterologous expression in *E. coli*, and *in vitro* testing of the purified protein in the presence of OME and UDP-glucose revealed the production of a compound matching the retention time, m/z and fragmentation pattern of the peak found in olive fruits (**Figure 5**.) To confirm the identity of this product, we performed a large-scale reaction, purified the resulting peak, and confirmed the structure via NMR, identifying the compound as 7-β-1-D-glucopyranosyl oleoside-11-methyl ester (**Supplementary Figures 1-7**). We thus named the enzyme *O. europaea* oleoside-11-methyl ester glucosyl transferase (OeOMEGT.)

**Figure 5.**
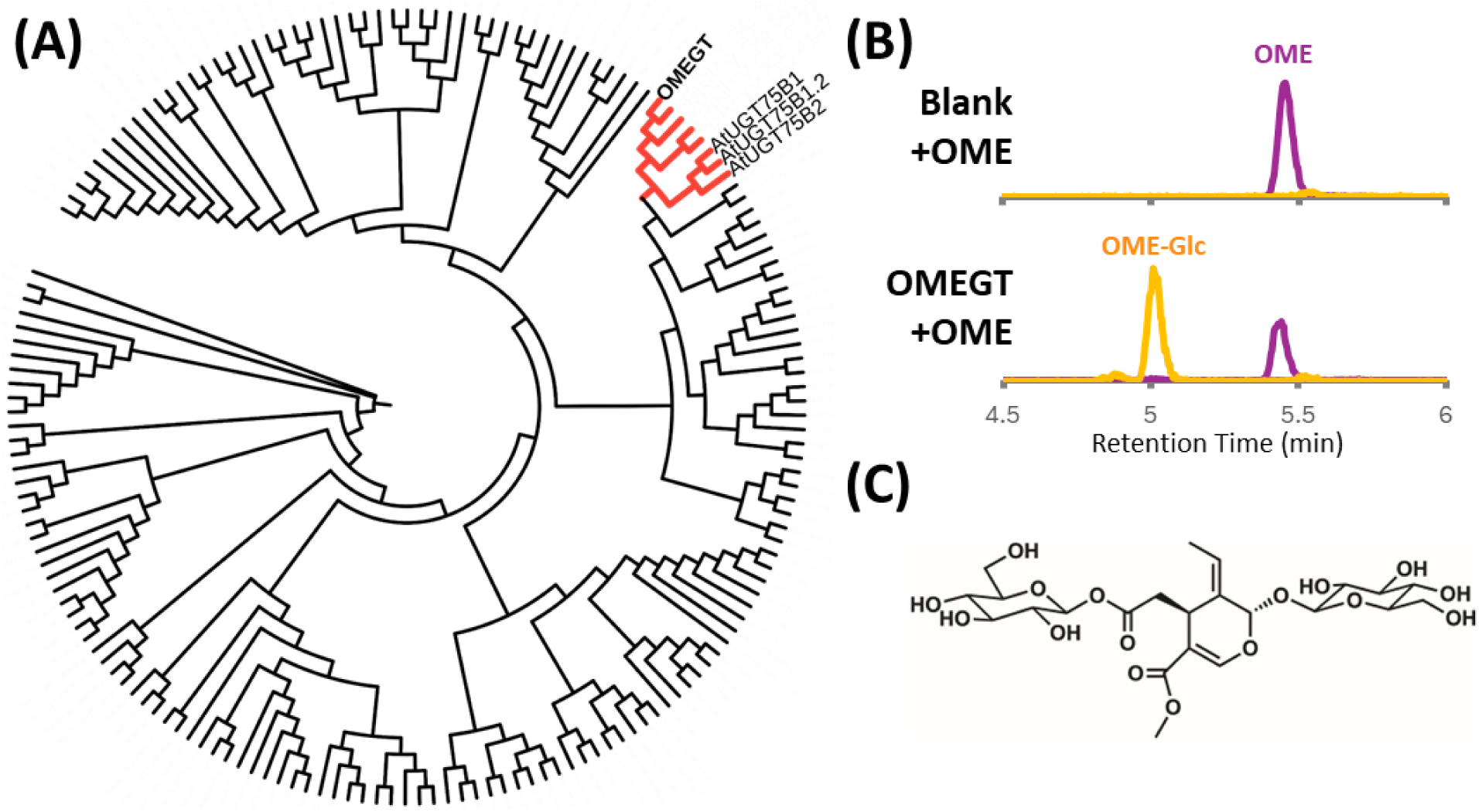
Assays of oleoside-11-methyl ester glucosyl transferase (OMEGT.) (A) Best-fit tree (WAG+F+R6) of a MUSCLE alignment of the protein sequences of *Olea europaea* and *Arabidopsis thaliana* UGTs. In red, we highlight the clade corresponding to the closest olive homologues of AtUGT75 proteins, with the position of the oleoside-11-methyl ester glucosyl transferase (OMEGT) discovered in this article. (B) Extracted ion chromatogram (XIC) of the most abundant oleoside methyl ester (OME; purple; [M-H]^−^ = 403.1246 ± 0.05) and its glucoside adduct (OME-Glc; orange; [M+formate]^−^ = 611.1829 ± 0.05) of the negative control reaction (Blank + OME; top) and the *in vitro* reaction with purified protein (OMEGT+OME; bottom.) (C) NMR confirmed structure of the HPLC-purified product of the reaction: 7-β-1-D-glucopyranosyl oleoside-11-methyl ester (OME-Glc.)

### Oleuropein synthase (OS)

Oleuropein is presumed to be produced by oxidation of the hydroxytyrosol moiety of ligstroside, as suggested by labelling experiments in *Syringa josikaea* (Damtoft, Franzyk, and Jensen 1993). It has been recently reported that olive polyphenol oxidases (PPOs) have the capacity to oxidize tyrosol, hydroxytyrosol and some of its esters (Derardja et al. 2024; Sánchez et al. 2023), but the products of the oxidation reactions of phenolic esters were not reported, and ligstroside was not included in the panel of substrates. We thus decided to narrow our search to the seven genes in our assembly that were differentially downregulated through ripening and were annotated as PPOs. When coinfiltrating *Nicotiana benthamiana* leaves with ligstroside and Agrobacterium harboring the transcripts TRINITY_GG_32073_c0_g1_i1 and TRINITY_GG_25161_c0_g1_i1, oleuropein was detected (**Figure 6**), and thus these sequences were named oleuropein synthase one (OeOS1) and two (OeOS2) respectively, based on expression levels, sharing only 46.5% amino acid identity. Noticeably, the sequence of OeOS2 has 99% identity to the enzyme OePPO3 reported by (Sánchez et al. 2023). We also tested a sequence (TRINITY_GG_32052_c0_g1_i1) with 79.8% amino acid identity to OeOS1, which was reported by Liu *et al*. (2023) to be syntenic with OeOS1, and probably a result of a recent duplication event. We found this sequence to have a detectable oleuropein synthase activity, thus naming it OeOS3 (**Figure 6**).

**Figure 6.**
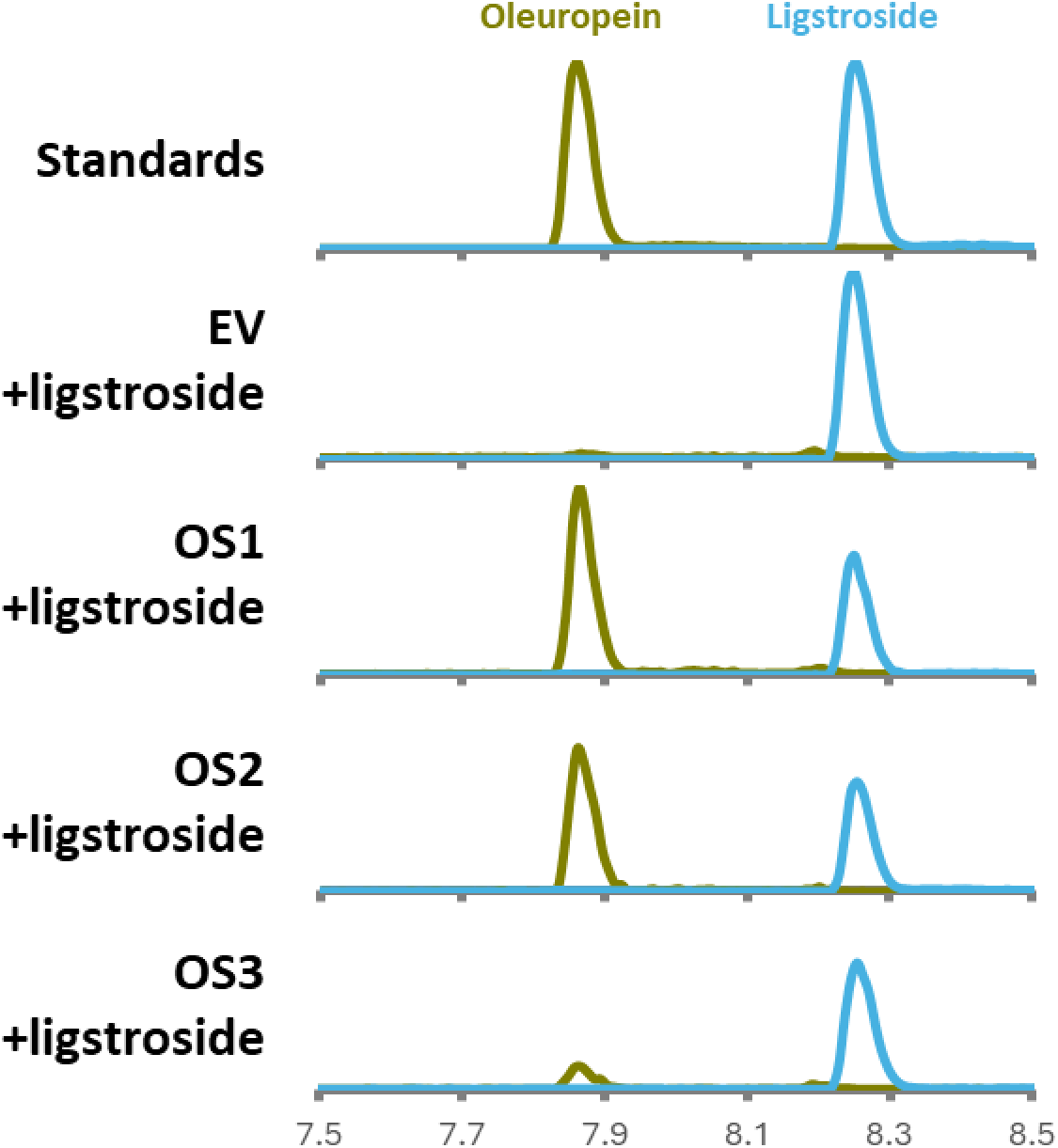
Extracted ion chromatograms (XIC) of co-infiltration assays. In olive green, the XIC of the most abundant oleuropein adduct ([M-H]^−^ = 539.1770 ± 0.05) is shown, and in cornflower blue, the most abundant ligstroside adduct ([M-H]^−^ = 523.1821 ± 0.05). From top to bottom: mix of standards, and leaf extracts of *Nicotiana benthamiana* co-infiltrated with ligstroside and Agrobacterium harboring either an empty vector (EV), oleuropein synthase one (OS1), two (OS2), or three (OS3).

### Pathway reconstitution

The enzymatic responsible for conversion of OME or OME-Glc to ligstroside was not identified, despite extensive screens of a wide variety of enzyme candidates. However, we could reconstitute the late-stage intermediate, 7-β-1-D-glucopyranosyl oleoside-11-methyl ester (OME-Glc), in *N. benthamiana* by transient, sequential expression of 7eLAS, 7eLAMT, OMES and OMEGT enzymes from olive, and feeding by the initial substrate, 7-deoxy-loganic acid (7-DLA). As it can be seen in **Figure 7**, upon 7-DLA infiltration, 7eLAS produces 7-*epi*-loganic acid, which is converted into 7-*epi*-loganin when 7eLAMT is added to the mixture; then OME is produced, via ketologanin, when OMES is included, and finally, OME-Glc is produced by addition of OMEGT, proving that the combination of these enzymes sequentially converts 7-deoxy-loganic acid to 7-β-1-D-glucopyranosyl oleoside-11-methyl ester (**Figure 7**.)

**Figure 7.**
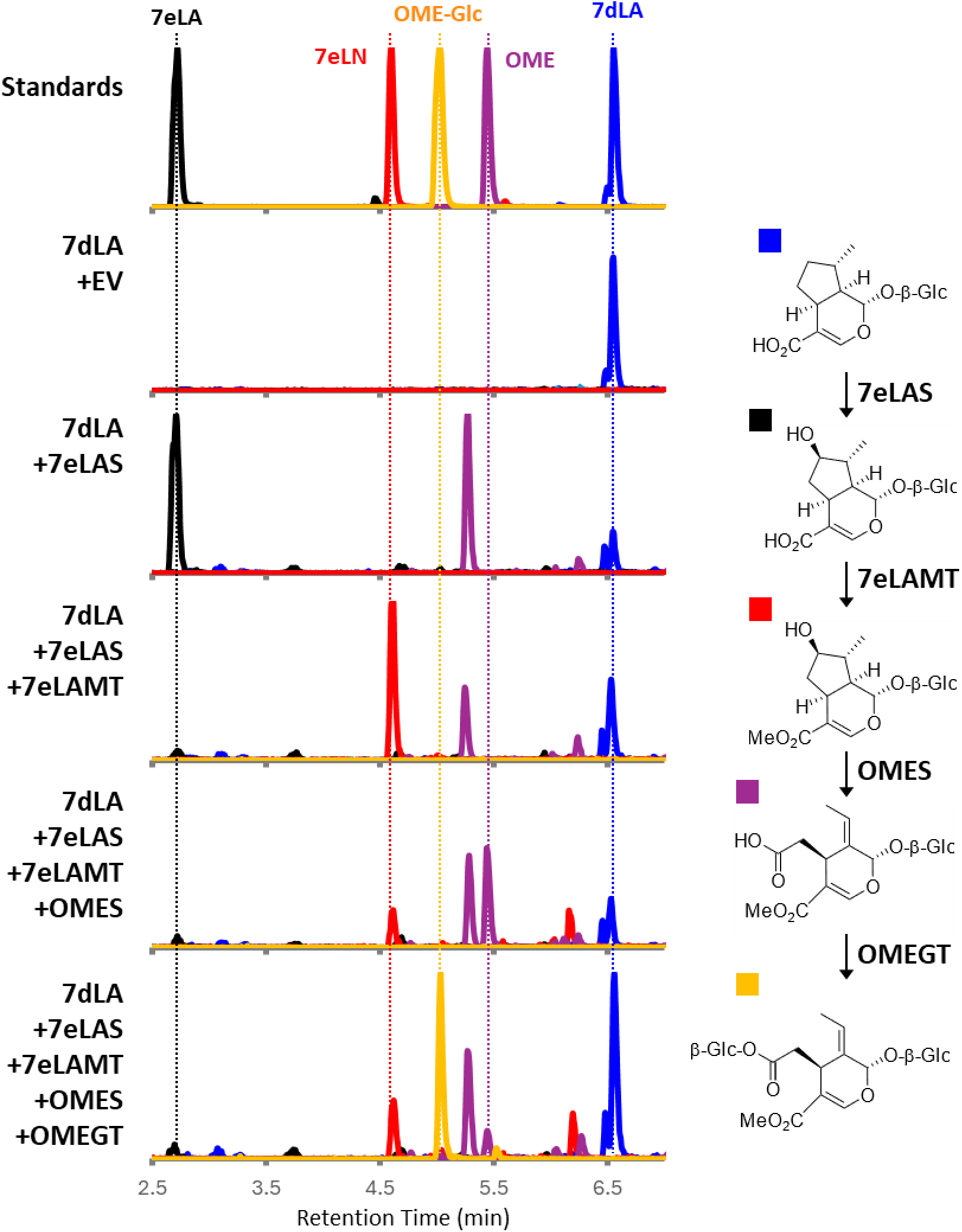
Pathway reconstruction in *Nicotiana benthamiana*. Extracted ion chromatograms of the most abundant adducts of 7-deoxy-loganic acid (blue, 7dLA, [M-H]^-^ = 359.1348 ± 0.05) 7-*epi*-loganic acid (black, LA, [M-H]^-^ = 375.1297 ± 0.05); 7-*epi*-loganin (red, 7eLN, [M+formate]^-^ = 435.1508 ± 0.05); oleoside methyl ester (purple, OME, [M-H]^−^ = 403.1246 ± 0.05) and 7-β-1-D-glucopyranosyl oleoside-11-methyl ester (orange, OME-Glc, [M+formate]^−^ = 611.1829 ± 0.05). From top to bottom: mix of standards (Standards), and extracts of *N. benthamiana* leaves infiltrated with 7-deoxy-loganic acid (7dLA) and Agrobacterium containing an empty vector (EV) and sequential coinfiltrations of 7-*epi*-loganic acid synthase (+7eLAS), 7-*epi*-loganic acid O-methyltransferase (+7eLAMT), oleoside methyl ester synthase (+OMES), and oleoside methyl ester glucosyl transferase (+OMEGT.) On the right, a schematic of the reconstructed biosynthetic module.

## DISCUSSION

Ripening of olive fruit is a complex process, involving change of color from green to dark purple, as well as lipid accumulation and organoleptic changes that make the fruit more palatable for seed dispersers. Our RNA-Seq results, consistent with previous reports (reviewed by Skodra et al. 2021), reflect the biology of the olive drupe as a sink tissue, as we see an increase in transcript levels from genes related to receiving carbon in the form of soluble sugars, and rerouting from conversion to starch into fatty acid biosynthesis as fruit ripens (**Supplementary Table 2**.) At the same time, we observe a decrease in expression of genes related to chlorophyll biosynthesis, a phenomenon that is partially responsible for color change from green to purple. Among these changes, the enrichment analysis suggests a shift in isoprenoid biosynthesis from the plastidial MEP pathway, decreasing as fruit matures, to the cytosolic MVA pathway, which increases during the same process **(Figure 2, Supplementary Table 3).** This is accompanied by a decrease in accumulation of iridoids, which are derived from the geraniol produced via the MEP pathway (**Figure 2**.) We found that not only is oleuropein accumulation regulated through decreasing expression of biosynthetic genes, from the upstream MEP pathway to the here discovered Oleuropein Synthases (OS1, OS2 and OS3), but also a decline in oleuropein across cultivars that is likely due to degradation. We expect this finding to encourage olive breeders to explore alternative strategies, targeting the degradation of oleuropein in fruit to improve the fruit quality of cultivars destined to olive oil production.

A common assumption in gene discovery is that similar biosynthetic pathways must be catalyzed by homologous enzymes, to the point where traced pathways are assumed to be identical in all species. Under this assumption, we discovered a 7-*epi*-loganic acid O-methyltransferase (7eLAMT), identified by homology to *C. roseus* LAMT, an OME glucosyl transferase (OMEGT), aided by homology to *Arabidopsis thaliana* group L UGTs, and two oleoside synthases (OS1 and OS2), annotated as polyphenol oxidases. However, when testing for homologs of *C. roseus* 7DLH and other transcripts annotated as cytochrome p450, we did not find the expected activity. Therefore, in this case, we used a comparative co-expression method that combines guilt by association approaches with sequence orthology inference, allowing to group correlation modules between different species. This method assumes that genes in the same biosynthetic pathway are co-regulated within each species, and allows to explore broader hypotheses regarding identity of missing enzymes, since it reduces the number of candidate genes to a subset that can be manually analyzed using biochemical logic. By analyzing expression data from *J. sambac* and *F. excelsior*, and comparing, via orthology, the transcripts that were co-expressed with the known biosynthetic gene homologues, we reduced the number of olive candidate genes from several thousands to a few dozens, allowing the discovery of 7eLAS.

With this method, we discovered a 2-oxoglutarate dependent dioxygenase that catalyzes the stereoselective oxidation of 7-deoxy-loganic acid to 7-*epi*-loganic acid, named 7eLAS. This reaction is analogous to the oxidation of 7-deoxy-loganic acid to loganic acid, which is catalyzed in *C. roseus* by a cytochrome p450 from the CYP72 family, 7-deoxy-loganic acid hydroxylase (7DLH). The discovery of an ODD that catalyzes the activation of the C-H bond at C7 in 7-deoxy-loganic acid opens the field for circumventing 7DLH, a known bottleneck in MIA production. Overall, given the increasing availability of expression data from species across the plant kingdom, this method can be readily applied to plant natural products with undiscovered pathways.

## ACKNOWLEDGEMENTS

CERL would like to acknowledge the contribution of the School of Engineering and Sciences, from the Tecnológico de Monterrey, for funding travel and lodging of his short research stay at the Max Planck Institute for Chemical Ecology. OC was supported by the short-term mobility program of CNR, Italy during her stay at SOC Dept. ICE. OC acknowledges financial support of the project ALIFUN (ARS01_00783) and of the PRIMA project BiomeNext (J63C21000100006), both funded by the Ministry of University and Research, Italy. The authors kindly thank Maritta Kunert and Matilde Florean, from the Department of Natural Products Biosynthesis of the Max Planck Institute for Chemical Ecology, for their help with Mass Spectrometry and reconstitution of the pathway in *Nicotiana benthamiana*. The authors acknowledge the help of Luciana Baldoni, CNR IBBR Italy, in olive germplasm selection and suggestions for the realization of the present work.

## AUTHOR CONTRIBUTIONS

OC, SOC, LC, and CERL designed experiments and wrote the manuscript; CERL, YJ, OC and MOK identified, cloned and characterized enzymes; BH and RMA synthesized standards and substrates; MOK, SH, OC and YN purified and determined the structure of OME-Glc; CERL, AG, FA and EF performed bioinformatics analysis; FP, MCV, OC and SM provided plant material and extracted RNA and metabolites for transcriptomics and metabolomics analysis; OC and MOK performed infiltrations and reconstituted the pathway in *Nicotiana benthamiana*. All authors reviewed the manuscript.

## DATA AVAILABILITY

Raw RNA-Seq data will be available upon acceptance in the EBI European Nucleotide Archive and the sequences of the enzymes discovered in this article will be available in GenBank upon publication.

