## Supplementary Figures for "It runs in the family: Discovery of enzymes in the oleuropein pathway in Olive (*Olea europaea*) by comparative transcriptomics"

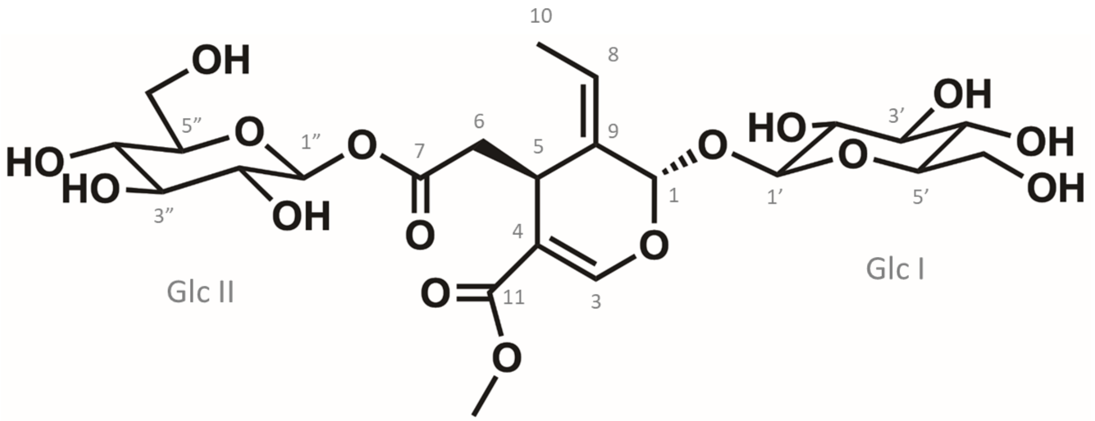


| pos. | $\delta_H$ | mult. | $J_{HH}$ | $\delta_C$ |
|---|---|---|---|---|
| 1 | 5.94 | *dd* | 1.3/1.1 | 95.2 |
| 3 | 7.53 | *s* | - | 155.2 |
| 4 | - | - | - | 109.0 |
| 5 | 4.01 | *dd* | 9.2/3.7 | 31.1 |
| 6a | 2.79 | *dd* | 15.4/3.7 | 40.4 |
| 6b | 2.59 | *dd* | 15.4/9.2 | 40.4 |
| 7 | - | - | - | 171.8 |
| 8 | 6.11 | *qd* | 7.1/1.1 | 125.3 |
| 9 | - | - | - | 130.0 |
| 10 | 1.77 | *dd* | 7.1/1.3 | 13.7 |
| 11 | - | - | - | 168.6 |
| OMe | 3.71 | *s* | - | 51.8 |
| **Glc I** | | | | |
| 1' | 4.81 | *d* | 7.9 | 100.7 |
| 2' | 3.31 | *m** | - | 74.7 |
| 3' | 3.41 | *dd* | 8.7/8.7 | 78.0 |
| 4' | 3.31 | *m** | - | 71.5 |
| 5' | 3.32 | *m** | - | 78.3 |
| 6'a | 3.90 | *bd* | 12.0 | 62.7 |
| 6'b | 3.67 | *dd* | 12.0/5.9 | 62.7 |
| **Glc II** | | | | |
| 1' | 5.43 | *d* | 8.2 | 95.7 |
| 2' | 3.33 | *dd* | 8.7/8.2 | 73.8 |
| 3' | 3.41 | *dd* | 9.1/8.7 | 78.0 |
| 4' | 3.35 | *m** | - | 71.0 |
| 5' | 3.35 | *m** | - | 78.6 |
| 6'a | 3.82 | *bd* | 12.3 | 62.3 |
| 6'b | 3.68 | *bd* | 12.3 | 62.3 |

* overlapped signals J unresolved

**Supplementary Figure 1. Chemical shift data for 7-β-1-D-glucopyranosyl oleoside-11-methyl ester.** Chemical shifts table (right) acquired at 700MHz in MeOH-$d_3$ is shown along with the elucidated structure.

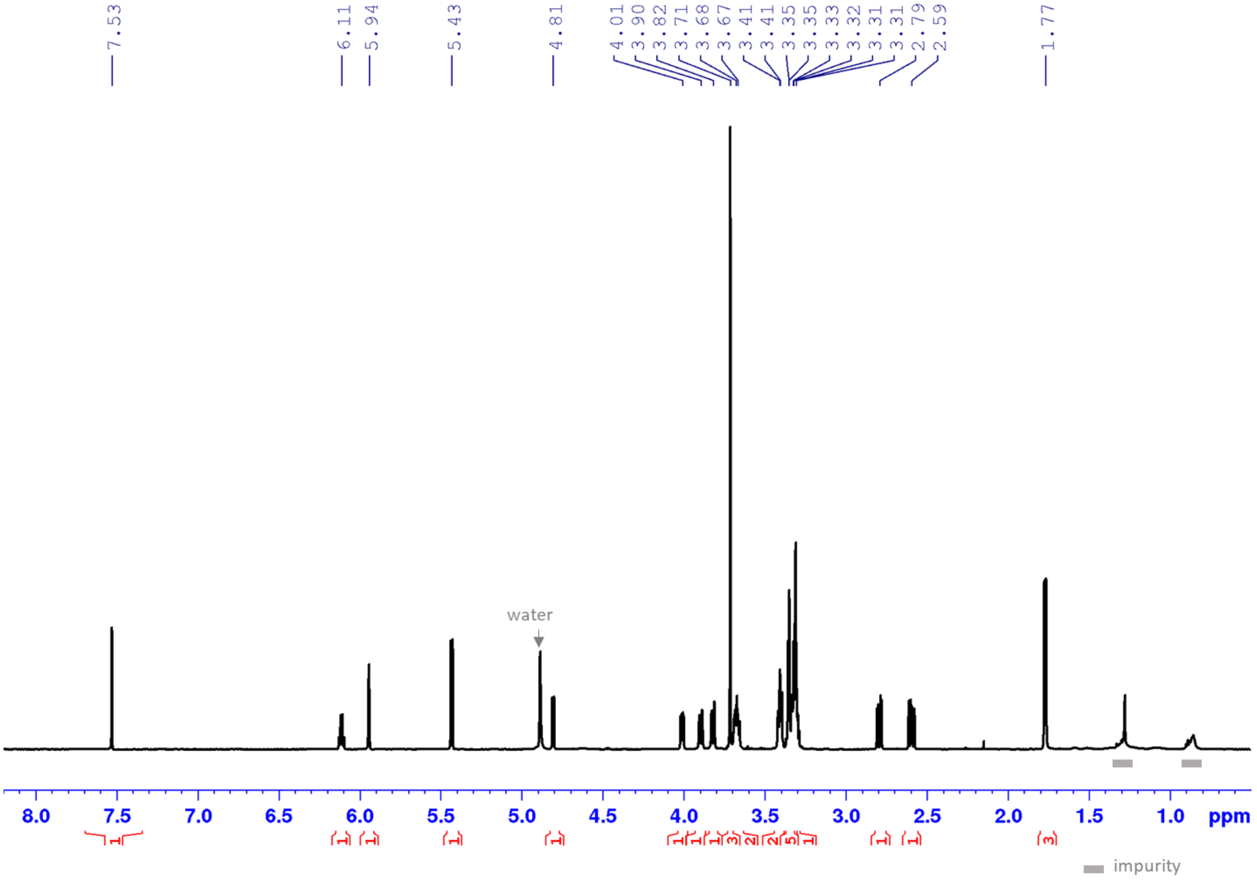


**Supplementary Figure 2. Proton NMR spectra for 7-β-1-D-glucopyranosyl oleoside-11-methyl ester with water suppression.** Full range in MeOH-*d*3 is shown.

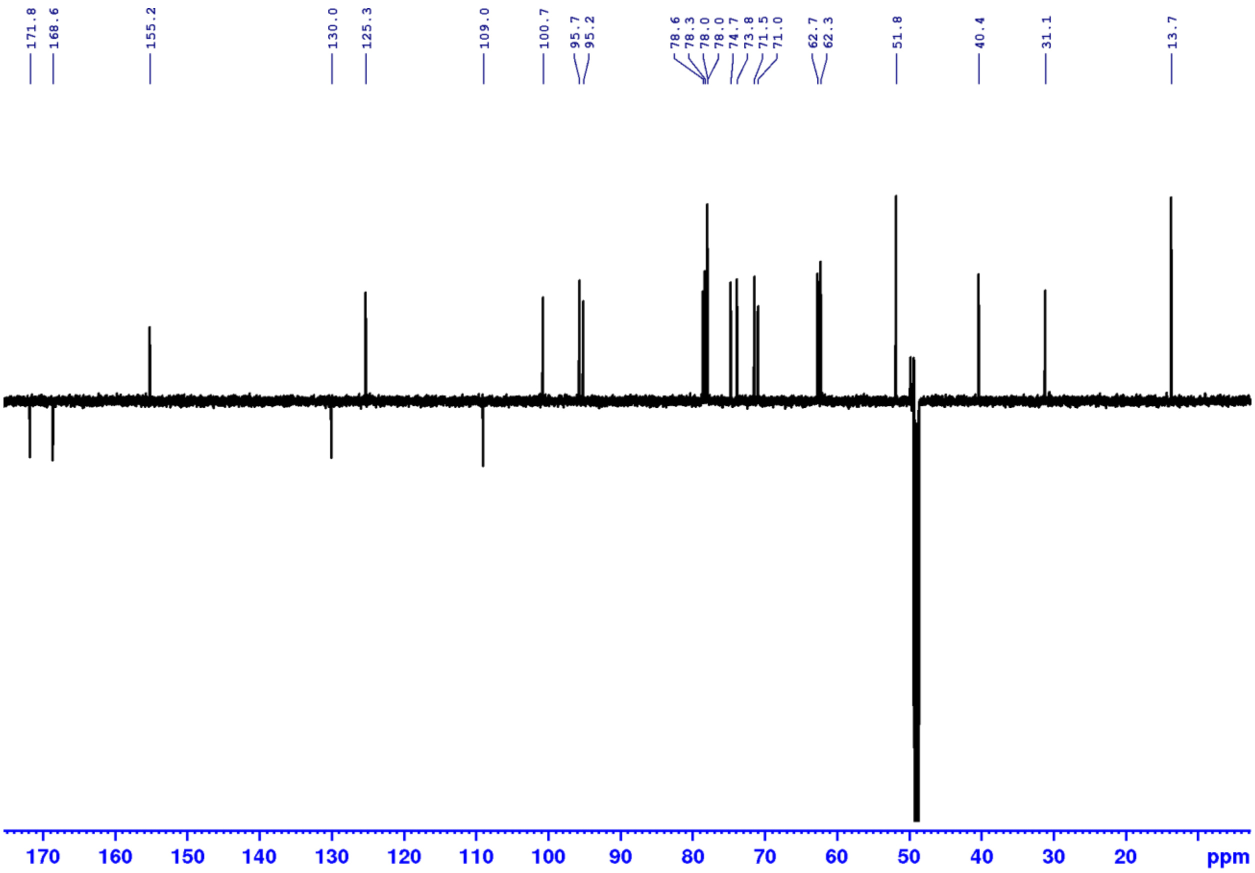


**Supplementary Figure 3. DEPTQ spectra for 7-β-1-D-glucopyranosyl oleoside-11-methyl ester.**

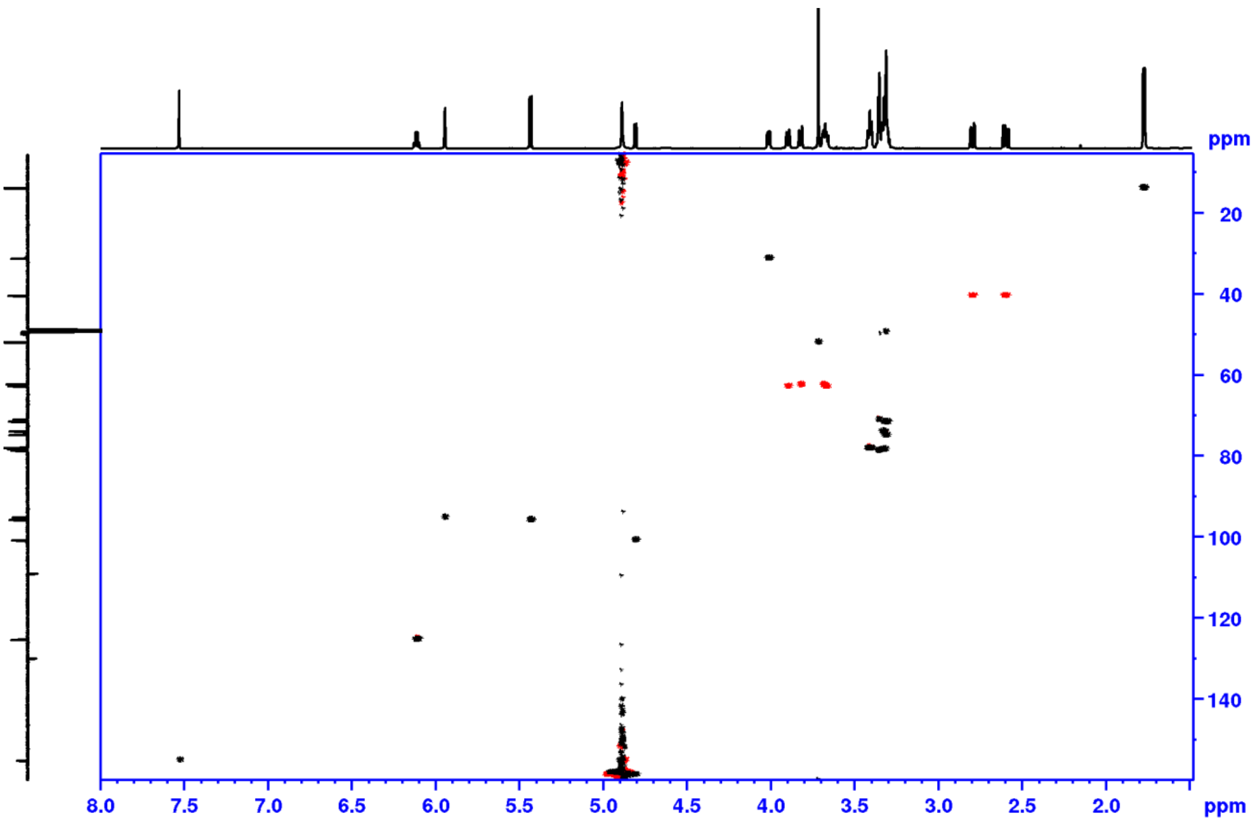


**Supplementary Figure 4. Full range phase sensitive HSQC spectra for 7-β-1-D-glucopyranosyl oleoside-11-methyl ester. $CH/CH_3$: black, $CH_2$: red.**

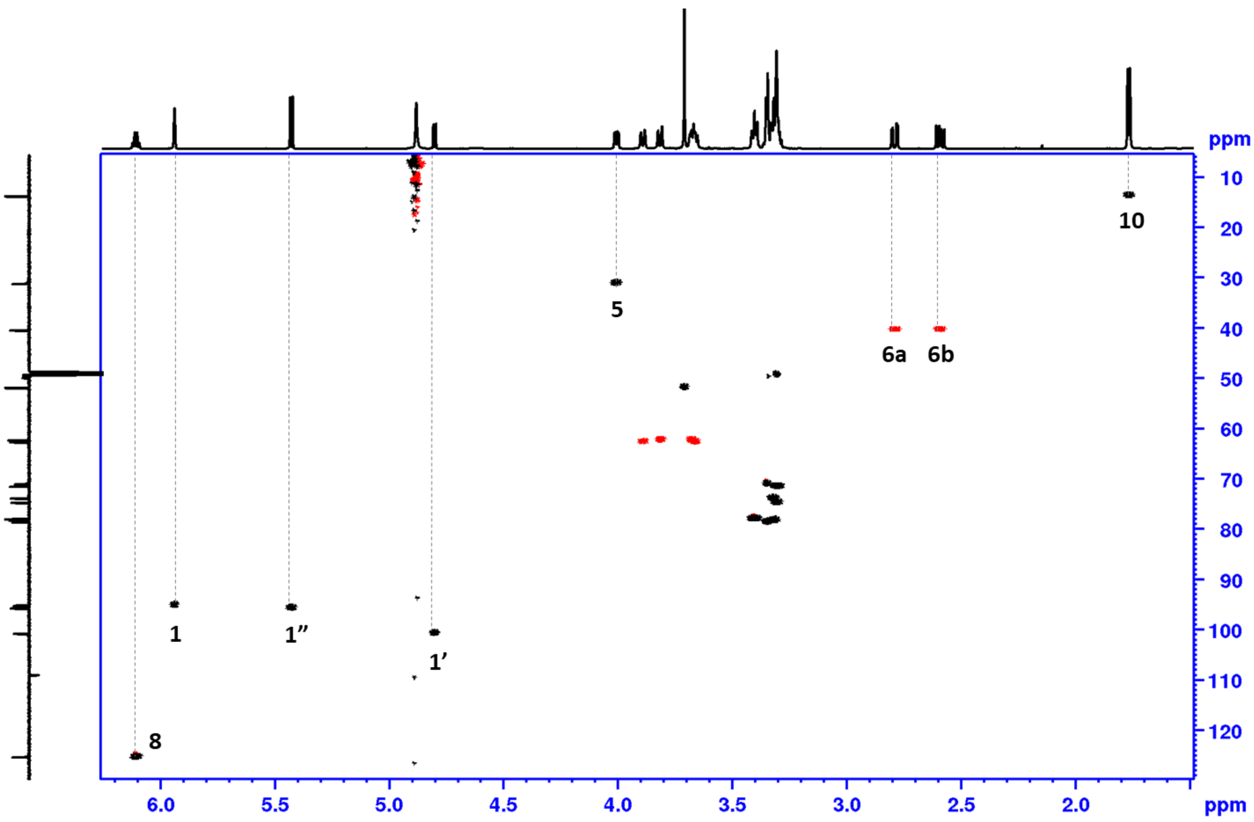


**Supplementary Figure 5. Focused phase sensitive HSQC spectra for 7-β-1-D-glucopyranosyl oleoside-11-methyl ester, from 1.5-6.5 ppm range. CH/$CH_3$: black, $CH_2$: red.**

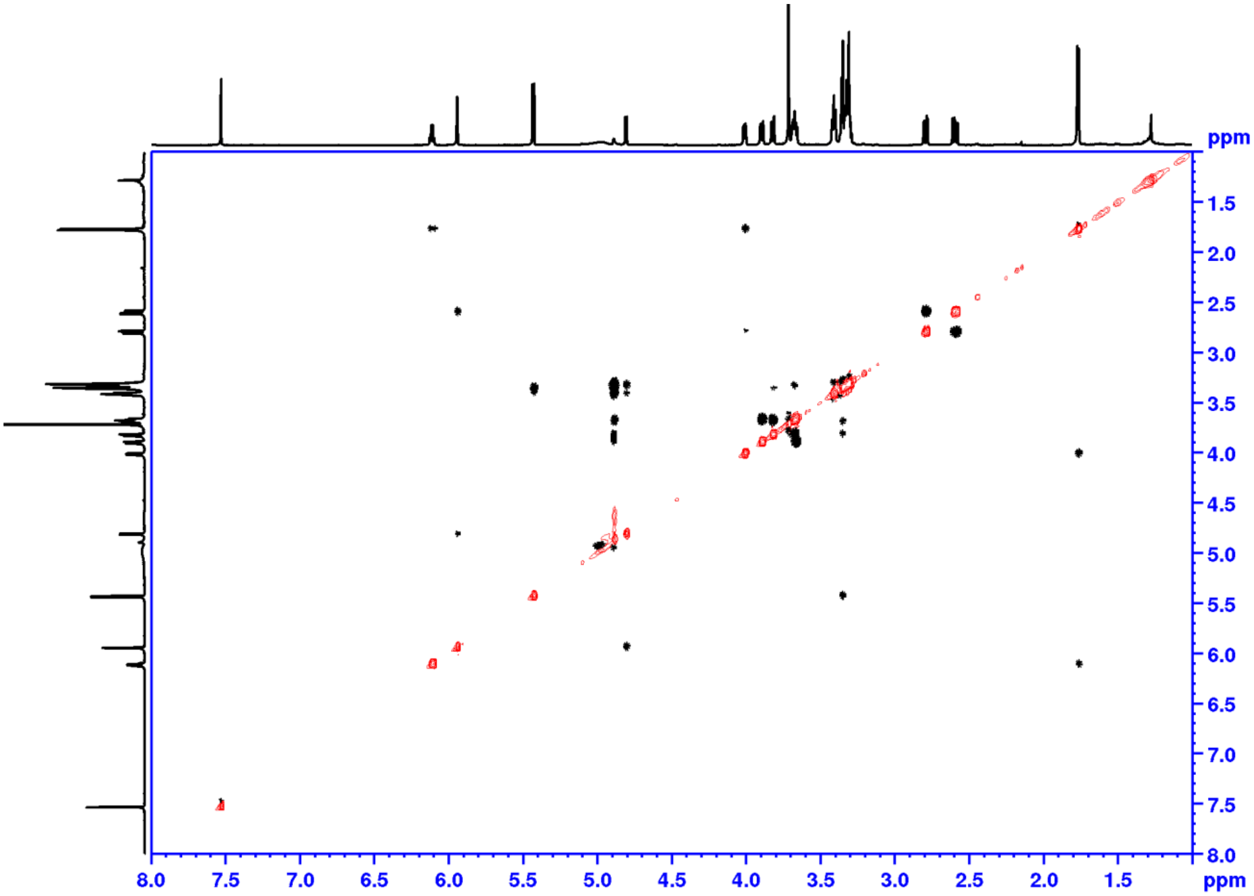


**Supplementary Figure 6. ROESY spectra for 7-β-1-D-glucopyranosyl oleoside-11-methyl ester, with water suppression.**

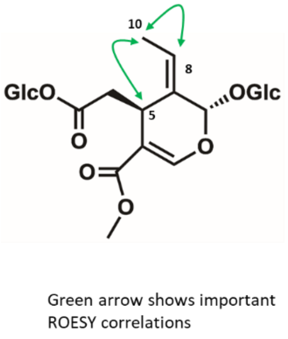


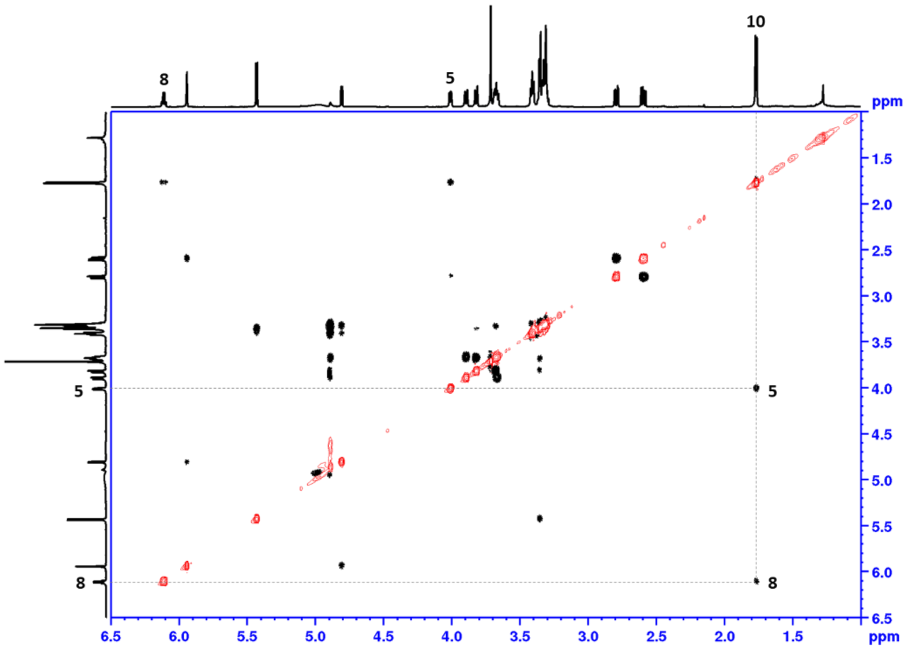


**Supplementary Figure 7. Focused ROESY spectra for 7-β-1-D-glucopyranosyl oleoside-11-methyl ester, with water suppression; range from 1.0-6.5 ppm.**

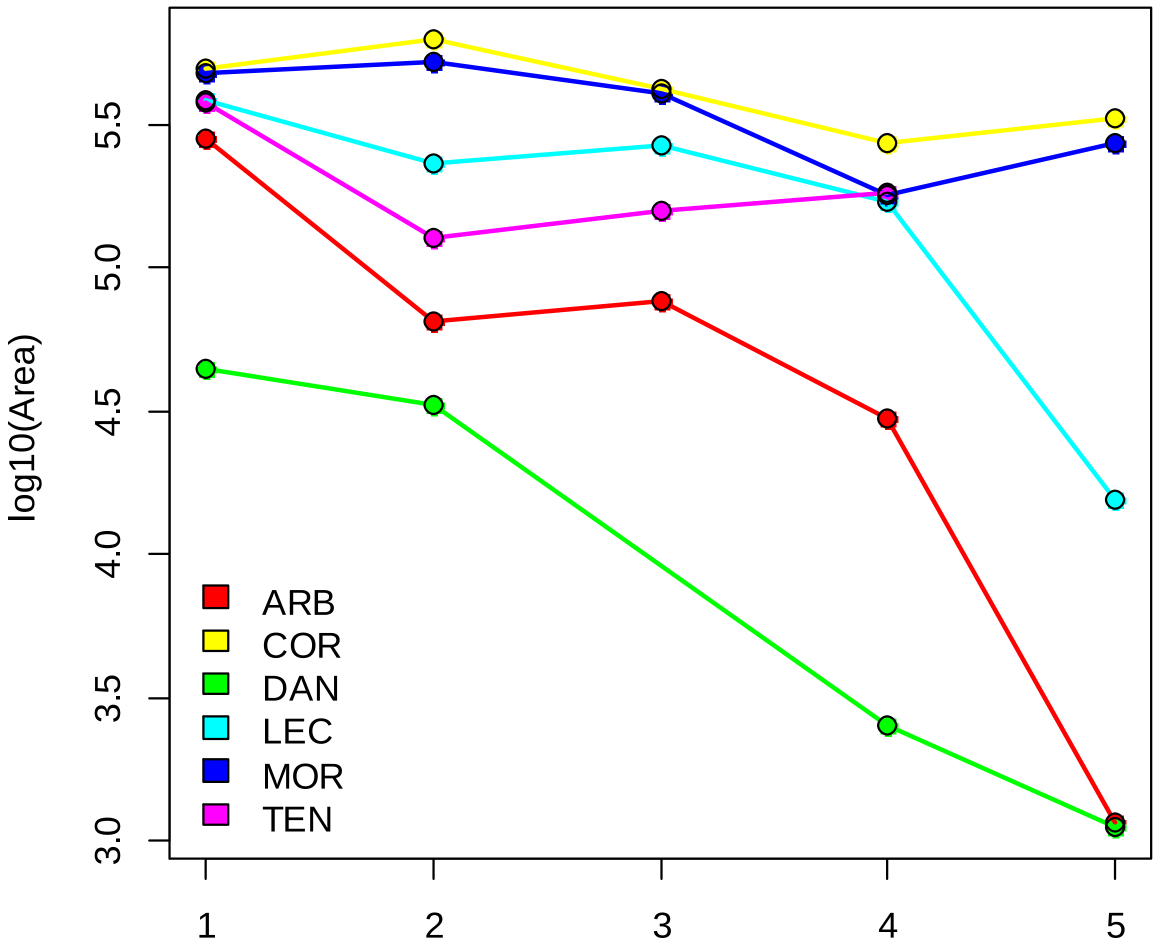


**Supplementary Figure 8. Oleuropein content of olive cultivars through time.** Logarithm of the area under the curve of the extracted ion chromatograms of the most abundant oleuropein adduct ($[M\text{-}H]^{-}$ = 539.1770 ± 0.05) of the cultivars *Dolce d'Andria* (DAN, green), *Tendellone* (TEN, purple)*, Arbequina* (ARB, red), *Leccino* (LEC, cyan), *Coratina* (COR, yellow) and *Moraiolo* (MOR, blue).

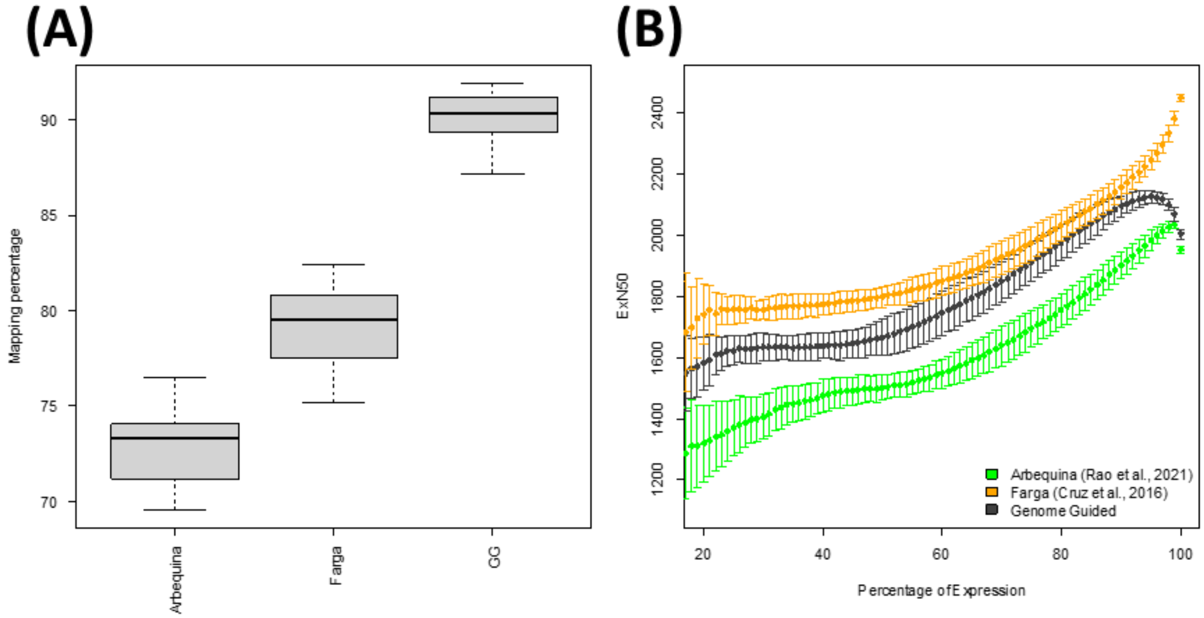


**Supplementary Figure 9. Assembly metrics.** (A) Mapping of the sequenced reads to the published Arbequina (Rao *et al.*, 2021) and Farga (Cruz *et al.*, 2016) genomes, as well as the genome guided assembly using Farga genome as a reference (GG.) (B) ExN50 of contigs, i.e. N50 as a function of percentage of expression of the top x-expressed genes of the mappings against Arbequina (green), Farga (yellow) and our genome guided assembly (black), expressed as mean values (solid dots) and standard deviation (error bars.)

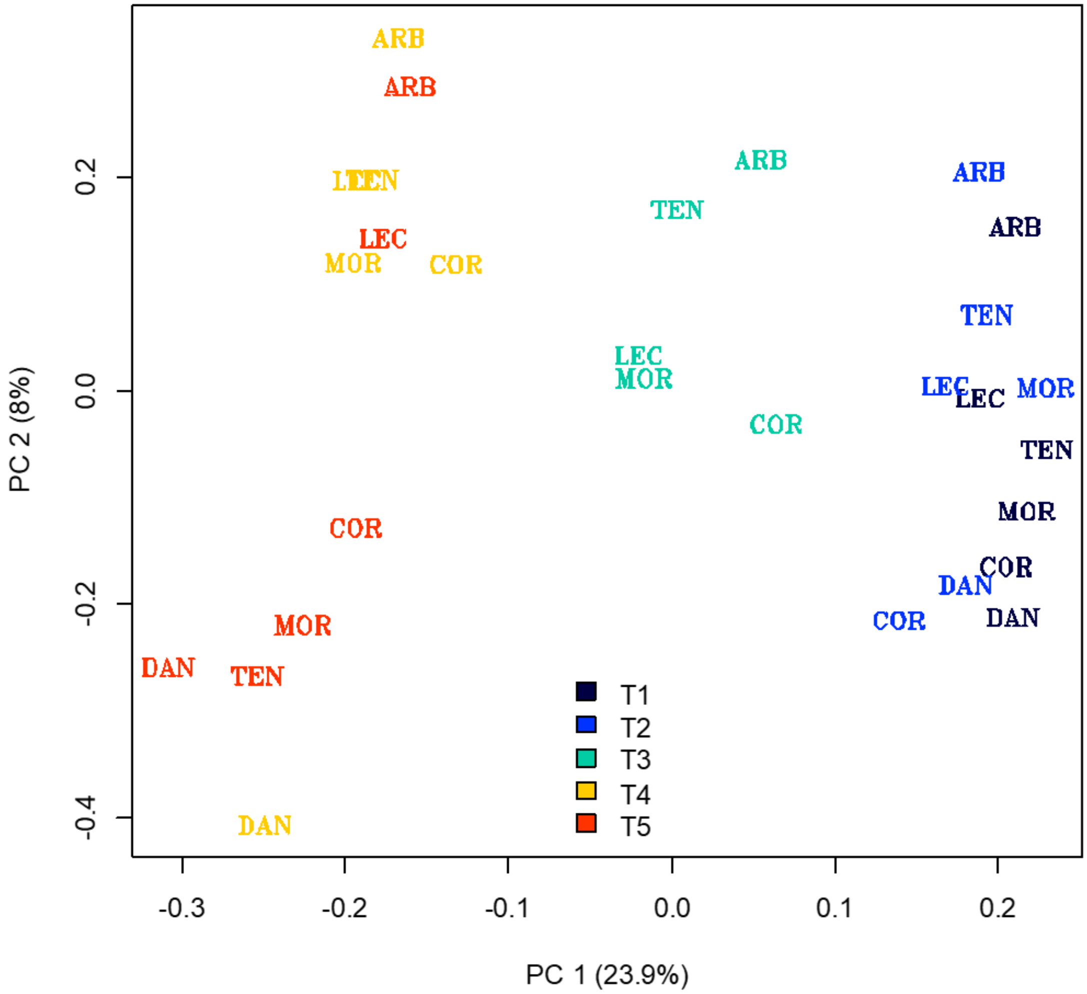


**Supplementary Figure 10. Principal Component Analysis (PCA) of expression data.** A loadings plot of the main two components of a PCA of the expression of all genes detected in samples through maturation. Labels correspond to cultivars *Dolce d'Andria* (DAN), *Tendellone* (TEN)*, Arbequina* (ARB), *Leccino* (LEC), *Coratina* (COR) and *Moraiolo* (MOR). Colors correspond to stages 1 (black), 2 (blue), 3 (cyan), 4 (yellow) and 5 (red).

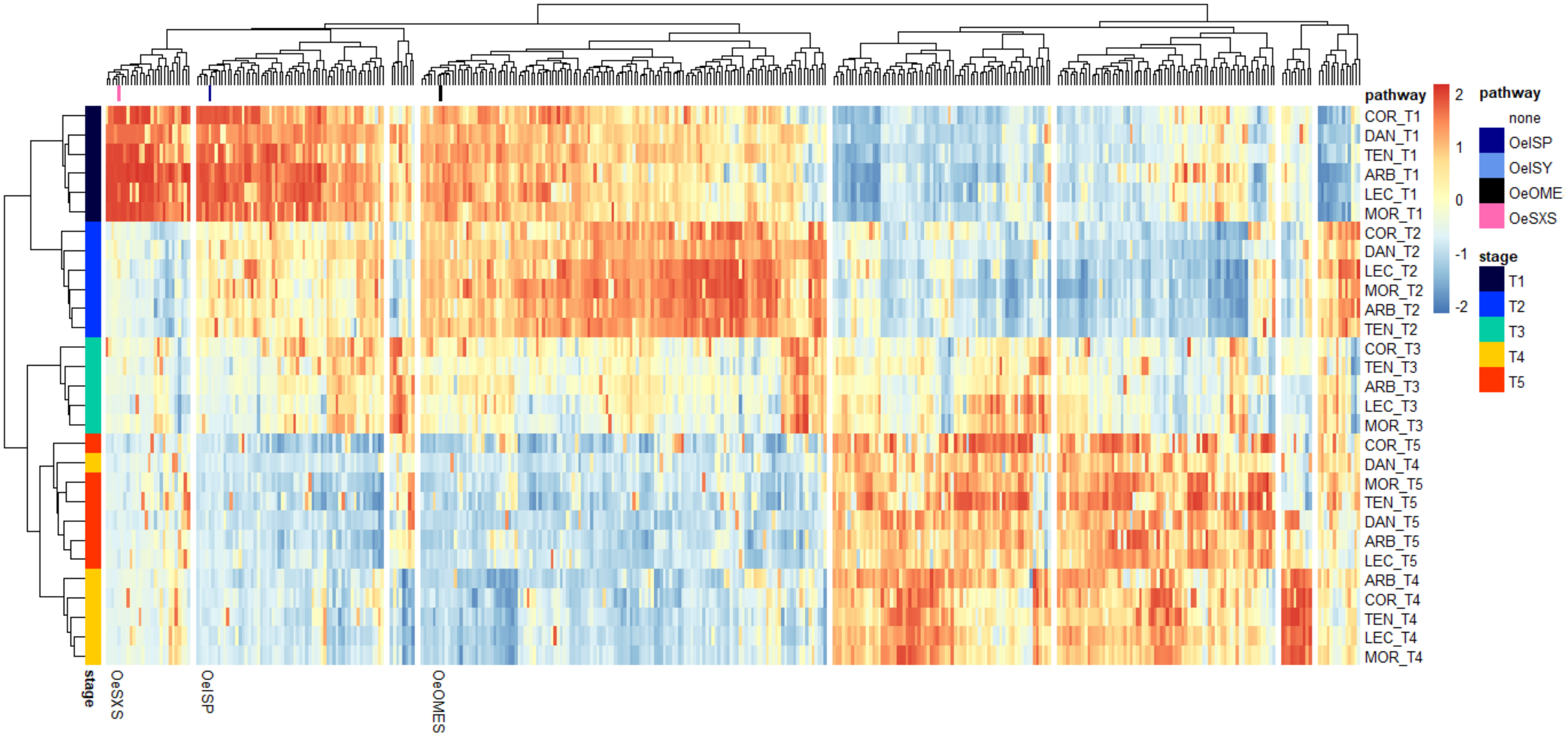


**Supplementary Figure 11. Heatmap of differentially expressed genes during Olive fruit maturation and ripening.** Heatmap of the codebook vectors of the 400 self-organizing map nodes, showing the type expression pattern of the 41,182 differentially expressed genes in olive fruit through maturation. Row band colors correspond to stages 1 (black), 2 (blue), 3 (cyan), 4 (yellow) and 5 (red).

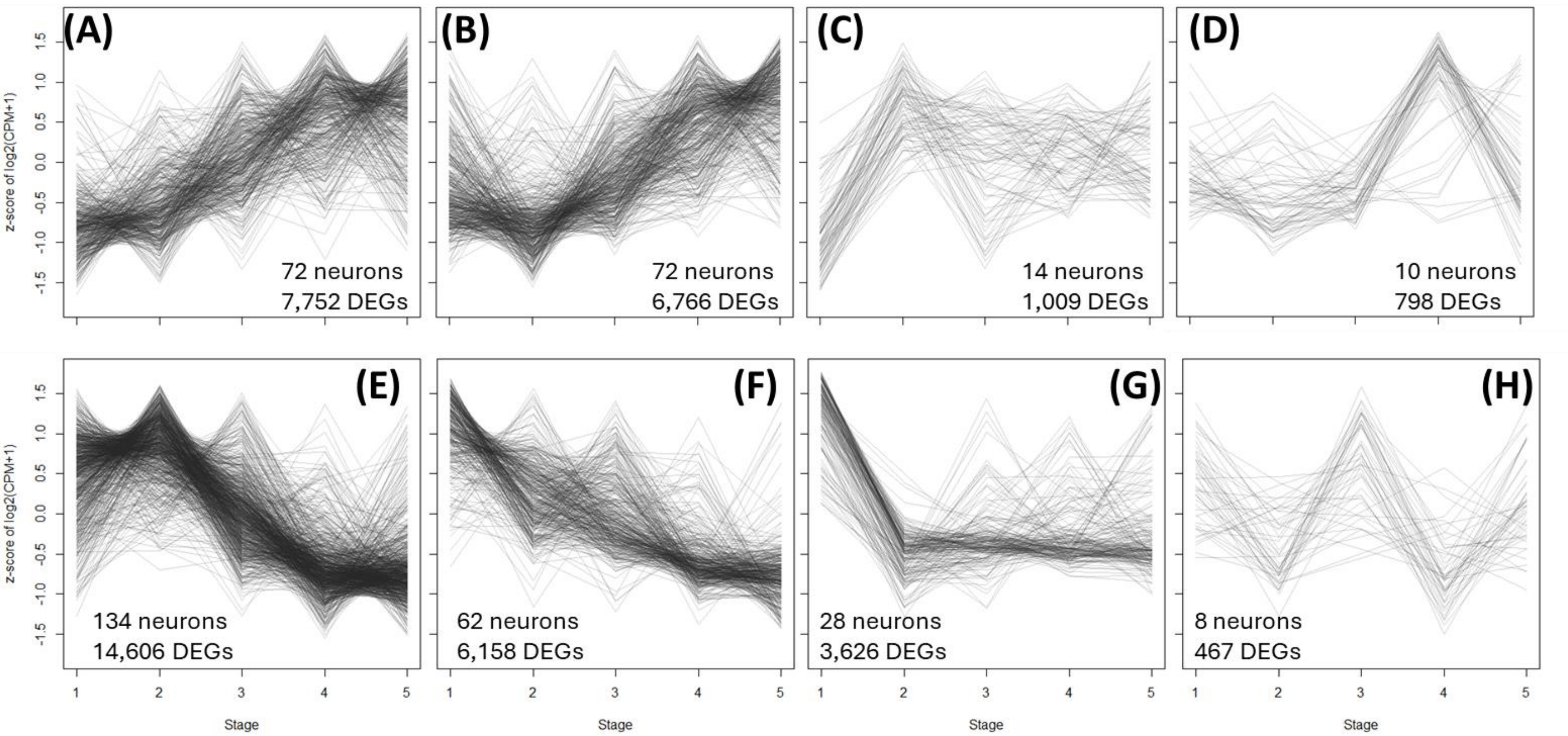


**Supplementary Figure 12. Type expression patterns of differentially expressed genes during Olive fruit maturation and ripening.** Line plots of the codebook vectors of the 400 self-organizing map nodes, grouped by cluster.

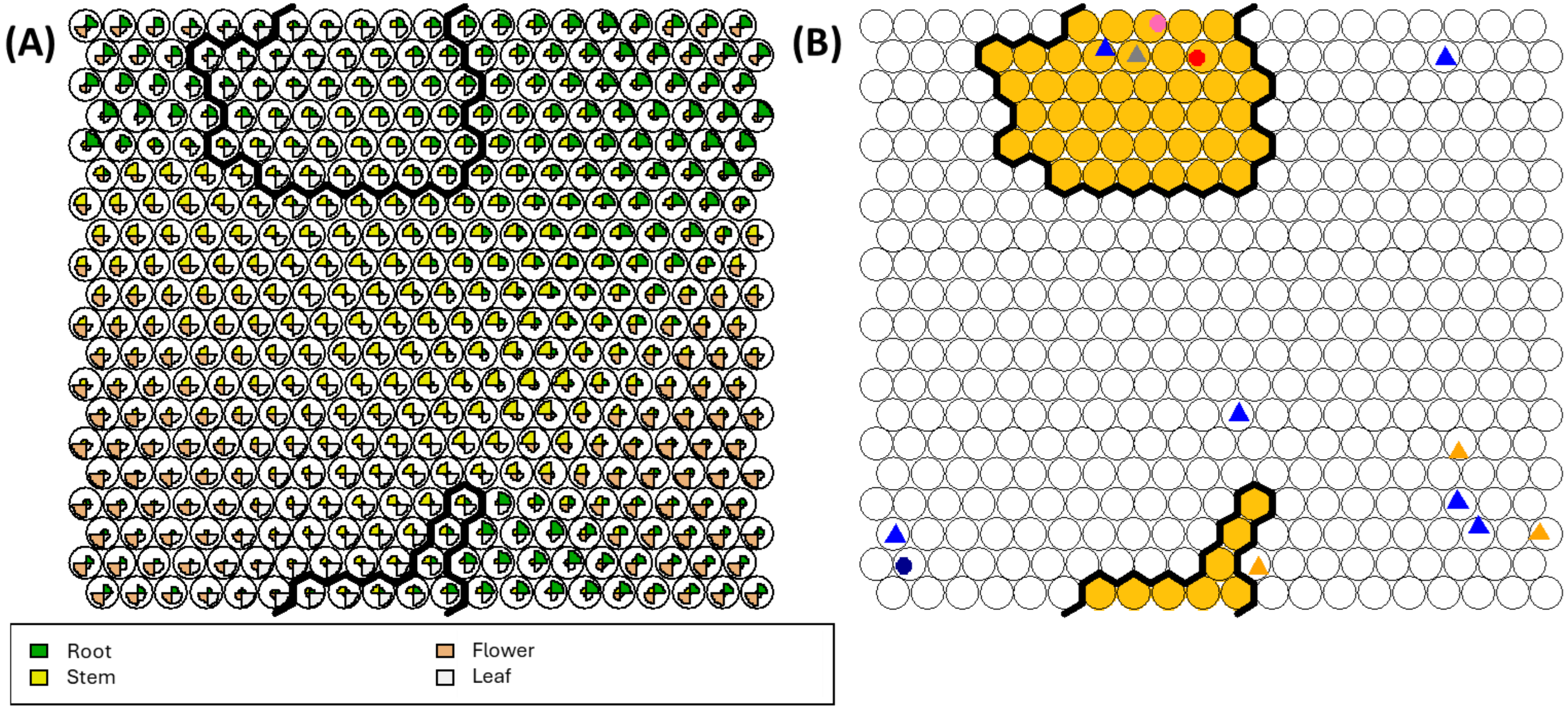


**Supplementary Figure 13. Self-Organizing Maps analysis of *Jasminium sambac*.** (A) Codes plot and (B) mapping plots of the self-organizing map summarizing expression data of *J. sambac*. The best BLAST results of known biosynthetic enzymes are shown in the codes plot as figures: ISY, iridoid synthase (black circle); ISP, iridoid synthase paralogue (gray triangle); IO, iridoid oxidase (pink circle); DLGT, 7-deoxyloganetic acid glucosyltransferase (blue triangle); 7eLAMT, 7-*epi*-loganic acid methyl-transferase (dark blue circle); OMES, oleoside methyl ester synthase (red circle); OMEGT, oleoside-11-methyl ester glucosyl transferase (orange triangle). In yellow, the selected cluster where most biosynthetic genes are located is highlighted.

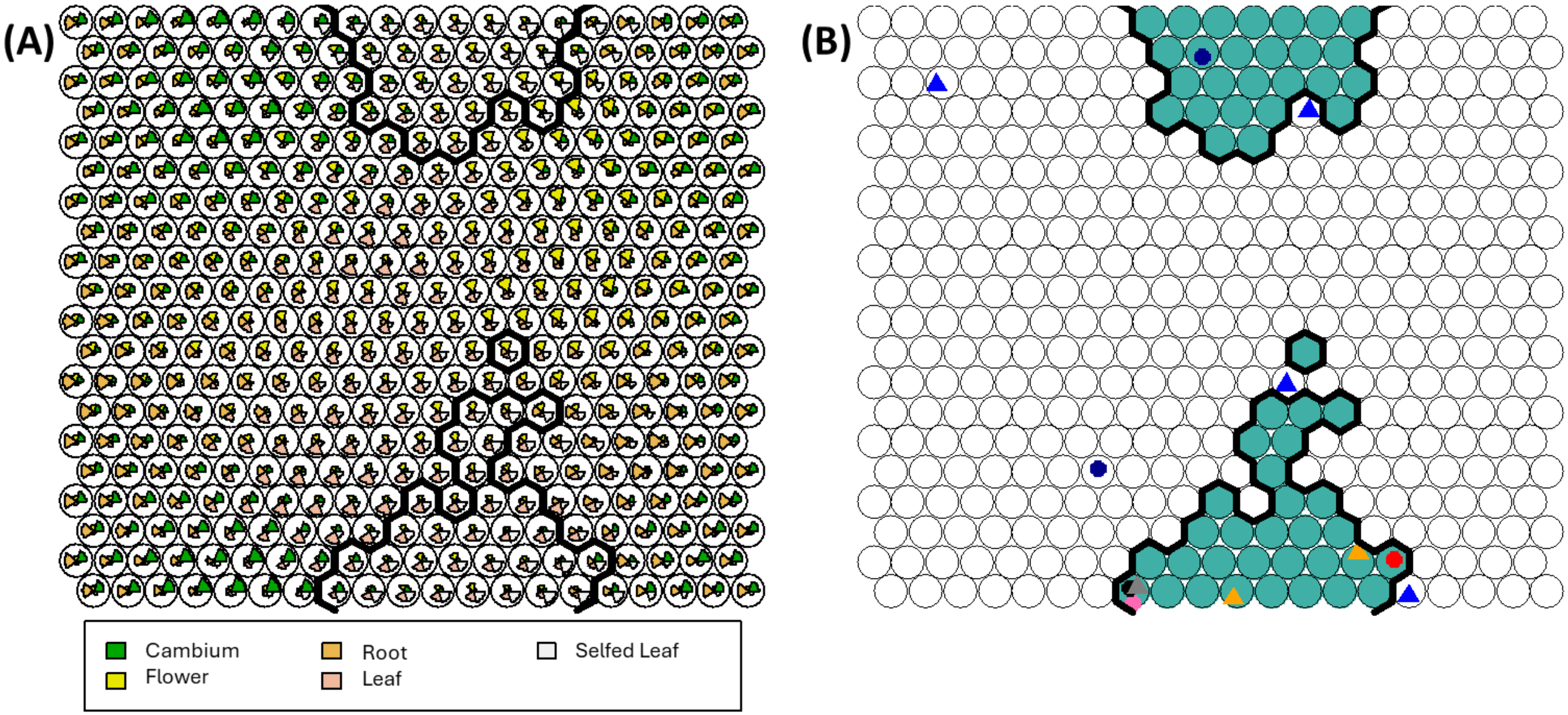


**Supplementary Figure 14. Self-Organizing Maps analysis of *Fracinus excelsior*.** (A) Codes plot and (B) mapping plots of the self-organizing map summarizing expression data of *F. excelsior*. The best BLAST results of known biosynthetic enzymes are shown in the codes plot as figures: ISY, iridoid synthase (black circle); ISP, iridoid synthase paralogue (gray triangle); IO, iridoid oxidase (pink circle); 7DLGT, 7-deoxyloganetic acid glucosyltransferase (blue triangle); 7eLAMT, 7-epi-loganic acid methyl-transferase (dark blue circle); OMES, oleoside methyl ester synthase (red circle); OMEGT, oleoside-11-methyl ester glucosyl transferase (orange triangle). In yellow, the selected cluster where most biosynthetic genes are located is highlighted.

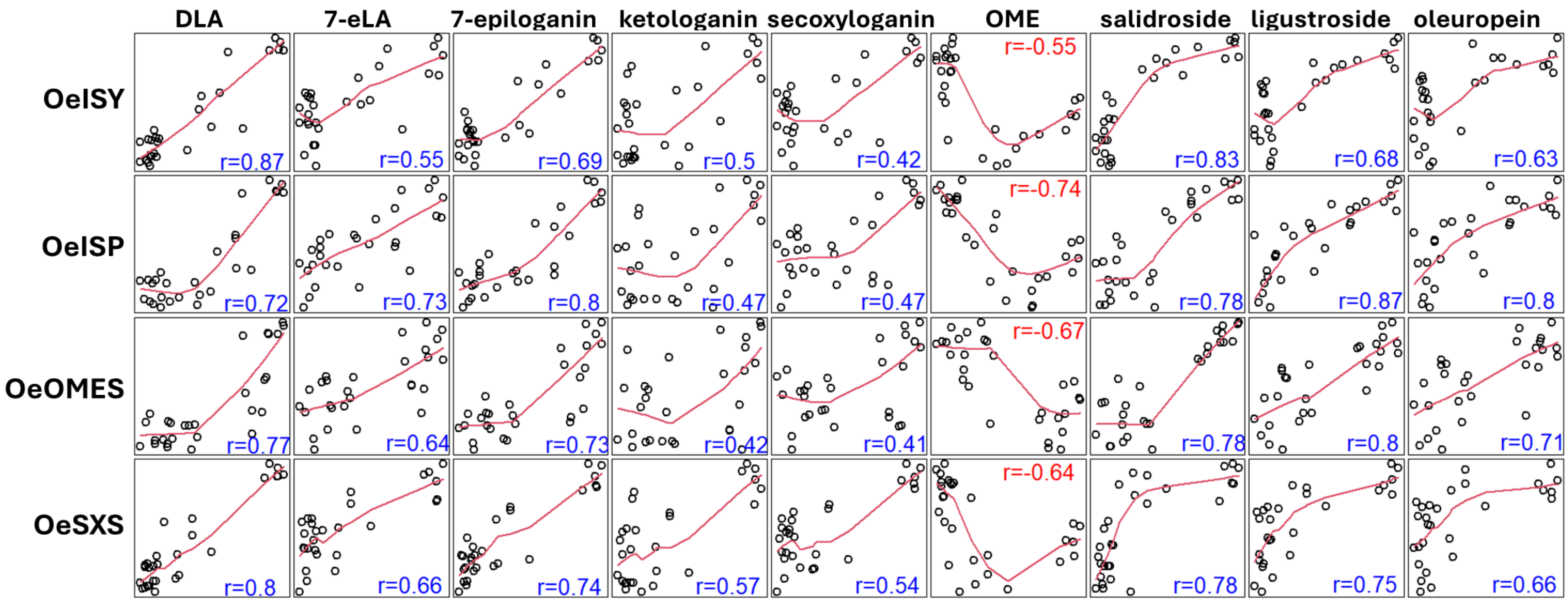


**Supplementary Figure 15. Correlations with known biosynthetic genes.** Scatter plots of center-scaled log-transformed areas of compounds (x-axis) and center-scaled expression data of known biosynthetic genes (y-axis). Positive (blue) and negative (red) Spearman correlations are shown for each gene-metabolite pair within the plot.
